# Visual recognition is heralded by shifts in local field potential oscillations and inhibitory networks in primary visual cortex

**DOI:** 10.1101/2020.12.22.423969

**Authors:** Dustin J. Hayden, Daniel P. Montgomery, Samuel F. Cooke, Mark F. Bear

## Abstract

Learning to recognize and filter familiar, irrelevant sensory stimuli eases the computational burden on the cerebral cortex. Inhibition is a candidate mechanism in this filtration process, and oscillations in the cortical local field potential (LFP) serve as markers of the engagement of different inhibitory neurons. We show here that LFP oscillatory activity in visual cortex is profoundly altered as male and female mice learn to recognize an oriented grating stimulus—low frequency (∼15 Hz peak) power sharply increases while high frequency (∼65 Hz peak) power decreases. These changes report recognition of the familiar pattern, as they disappear when the stimulus is rotated to a novel orientation. Two-photon imaging of neuronal activity reveals that parvalbumin-expressing inhibitory neurons disengage with familiar stimuli and reactivate to novelty, whereas somatostatin-expressing inhibitory neurons show opposing activity patterns. We propose a model in which the balance of two interacting interneuron circuits shifts as novel stimuli become familiar.

**Significance Statement:** Habituation, familiarity and novelty detection are fundamental cognitive processes that enable organisms to adaptively filter meaningless stimuli and focus attention on potentially important elements of their environment. We have shown that this process can be studied fruitfully in the mouse primary visual cortex by using simple grating stimuli for which novelty and familiarity are defined by orientation, and by measuring stimulus-evoked and continuous local field potentials. Altered event-related and spontaneous potentials, and deficient habituation, are well-documented features of several neurodevelopmental psychiatric disorders. The paradigm described here will be valuable to interrogate the origins of these signals and the meaning of their disruption more deeply.

## Introduction

The awake brain receives a steady stream of sensory stimuli. Distinguishing novel, potentially relevant stimuli from familiar, irrelevant stimuli is essential for the dedication of energy and attention to only those elements of the environment that may be salient for survival. Previous studies have described an electrophysiological signature of long-term recognition memory within primary visual cortex (V1) of mice that is highly selective for stimulus attributes, such as orientation (Frenkel et al., 2006; Cooke and Bear, 2010). Over days of repeated presentation of a simple, phase-reversing sinusoidal grating stimulus, the magnitude of visually-evoked potentials (VEPs) recorded in layer 4 of V1 in awake, head-fixed mice significantly increases. We refer to this process as stimulus-selective response plasticity (SRP). Similar phenomena have been reported by others (Aton et al., 2014; Kaneko and Stryker, 2014; Kaneko et al., 2017; Kissinger et al., 2018).

Much headway has been made in understanding the requirements for SRP. Disruption of SRP by treatments local to V1, notably including manipulations of NMDA receptor (NMDAR) function and AMPA receptor trafficking in principal cells, had suggested involvement of the mechanisms of long-term potentiation (LTP) of feedforward excitatory synapses (Frenkel et al., 2006; Cooke et al., 2015). However, more recent studies show that SRP (1) is not supported by plasticity at excitatory layer 4 synapses (Cooke and Bear, 2014; Fong et al., 2020) and (2) depends on the activity of parvalbumin-expressing (PV+) neurons in V1 (Kaplan et al., 2016). Given the extensive evidence that PV+ inhibitory neurons contribute to “gamma” oscillations (≥ 40 Hz) in the cortex (Cardin et al., 2009; Korotkova et al., 2010; Carlen et al., 2012; Gonzalez-Burgos and Lewis, 2012; Lewis et al., 2012; Kuki et al., 2015; Jadi et al., 2016; Polepalli et al., 2017), in the current study we sought to understand how the local field potential (LFP) and PV+ cell activity in layer 4 are influenced by stimulus familiarity and how these changes evolve over time.

We found that exposure of mice to a novel visual stimulus elicits high frequency (∼65 Hz) oscillations in the LFP and increases PV+ cell activity measured using two-photon (2-p) calcium imaging. With repeated viewing over days, the now-familiar stimulus elicited reduced high frequency power in the LFP with a corresponding decrease in PV+ cell activity, as well as a sharp increase in low frequency (∼15 Hz, “beta”) oscillations. The observed increase in oscillations at this frequency is consistent with the hypothesis that familiar stimuli recruit a population of somatostatin-expressing (SOM+) interneurons (Kuki et al., 2015), as has been observed in auditory cortex following long-term habituation with passive sound exposure (Kato et al., 2015). Indeed, 2-p calcium imaging in SOM+ cells in V1 revealed an increase in their activity during familiar stimulus viewing. These signatures of familiar stimulus recognition did not appear immediately on stimulus onset, but developed rapidly within the first few seconds of visual stimulation. Our findings significantly advance the understanding of how visual recognition memory is expressed at the circuit level and demonstrate how stimulus familiarity imposes oscillatory activity on cerebral cortex.

## Materials and Methods

### Mice

All procedures adhered to the guidelines of the National Institutes of Health and were approved by the Committee on Animal Care at MIT, Cambridge, MA, USA. For local field potential experiments, we used male and female mice on a C57BL/6 background (Charles River laboratory international, Wilmington, MA). For calcium imaging experiments, we used male and female PV-Cre mice (B6.129P2-Pvalb^tm1(cre)Arbr^/J, RRID:IMSR JAX:017320) and SOM-Cre mice (B6N.Cg-Sst^tm2.1(cre)Zjh^/J, RRID:IMSR JAX:018973). The familiar-novel differences reported in this study did not differ qualitatively by sex so both were combined, in agreement with previous studies (Fong et al., 2020). Animals were housed in groups of 2-5 same-sex littermates after weaning at P21. They had access to food and water *ad libitum* and were maintained on a 12-hour light-dark cycle.

### Surgery

For local field potential experiments, young adult C57BL/6 mice (P26-P52) were first injected with 0.1 mg/kg Buprenex sub-cutaneously (s.c.) to provide analgesia. Induction of anesthesia was achieved via inhalation of isoflurane (3% in oxygen) and thereafter maintained via inhalant isoflurane (∼1-2% in oxygen). Prior to surgical incision, the head was shaved and the scalp cleaned with povidone–iodine (10% w/v) and ethanol (70% v/v). Scalp was resected and the skull surface was scored. A steel headpost was affixed to the skull (anterior to bregma) with cyanoacrylate glue. Small burr holes were drilled above both hemispheres of binocular V1 (3.0 mm lateral of lambda). Tapered 300-500 kΩ tungsten recording electrodes (FHC, Bowdoin, ME, US), 75 µm in diameter at their widest point were implanted in each hemisphere, 450 µm below cortical surface. Silver wire (A-M systems, Sequim, WA, US) reference electrodes were placed over the left frontal cortex. Electrodes were secured using cyanoacrylate and the skull was covered with dental cement. Nonsteroidal anti-inflammatory drugs were administered upon return to the home cage (meloxicam, 1 mg/kg s.c.). Signs of infection and discomfort were carefully monitored. Mice were allowed to recover for at least 48 hours prior to head-fixation.

For cranial window implantations for two-photon calcium imaging, adult PV-Cre or SOM-Cre mice (P43-P133) were anesthetized and prepared as described above. Following scalp incision, a lidocaine (1%) solution was applied onto the periosteum and the exposed area of skull gently scraped with a scalpel blade. Then, a 3 mm craniotomy was made over binocular V1. Adeno-associated virus containing the GCaMP7f gene (pGP-AAV9-syn-FLEX-jGCaMP7f-WPRE, 104488-AAV9, Addgene, Watertown, MA, USA) was loaded into a glass micropipette with a tip diameter of 40–50 µm attached to a Nanoject II injection system (Drummond Scientific, Broomall, PA, USA). The micropipette was then inserted into binocular V1 layer 4 at depths of 400 and 450 µm below pial surface and ∼50 nl of virus was delivered at each depth. Next, a sterile 3 mm round glass coverslip (CS-3R-0, Warner Instruments, Hamden, CT, USA) was gently laid on top of the exposed dura mater. The coverslip was secured with cyanoacrylate glue and a stainless steel head post was attached to the skull. Once the glue had set, dental acrylic (C&B Metabond Quick Adhesive Cement System, Parkell, NY, USA) was mixed and applied throughout the exposed skull surface.

### Visual stimulus delivery

Prior to stimulus delivery, mice were acclimated to head restraint in front of a gray screen for a 30-minute session on each of two consecutive days. After habituation, for the LFP, pupil, and movement experiments, mice were presented with 5 blocks of 100 phase-reversals of an oriented grating stimulus phase-reversing at 0.5 Hz. They were shown this stimulus for 6 consecutive days. On day 7, they were shown both the familiar stimulus orientation as well as blocks of a novel stimulus offset 90° from the novel orientation. Each stimulus block was preceded by a period of gray screen, a period of black screen, and another period of gray screen. Gray periods lasted 6 or 12 s and black periods lasted 10 or 20 s, depending on the recording system. Discrete sections of gray and black screen viewing were timestamped for later normalization. After habituation for the calcium imaging experiments, mice were presented with 5 blocks of 120 phase-reversals of an oriented grating stimulus phase-reversing at 0.5 Hz. They were shown this stimulus for 4 consecutive days. On day 5, they were shown both the familiar stimulus orientation as well as blocks of a novel stimulus offset 90° from the novel orientation. Each stimulus block was preceded by 30 seconds of gray screen. To keep head-restraint to a minimum during calcium imaging experiments, only 4 blocks of each stimulus were used on day 5. For all experiments, if more than 1 orientation was shown within a session, stimulus blocks were pseudo-randomly interleaved such that 3 consecutive presentations of the same stimulus never occurred. Visual stimuli consisted of full-field, 100% contrast, sinusoidal gratings that were presented on a computer monitor. Visual stimuli were generated using custom software written in either C++ for interaction with a VSG2/2 card (Cambridge Research systems, Kent, U.K.) or Matlab (MathWorks, Natick, MA, U.S.) using the PsychToolbox extension (http://psychtoolbox.org) to control stimulus drawing and timing. Grating stimuli spanned the full range of monitor display values between black and white, with gamma-correction to ensure constant total luminance in both gray-screen and patterned stimulus conditions.

### In vivo electrophysiology experimental design and analysis

Electrophysiological recordings were conducted in awake, head-restrained mice. Recordings were amplified and digitized using the Recorder-64 system (Plexon Inc., Dallas, TX, US) or the RHD Recording system (Intan Technologies, Los Angeles, CA, US). Two recording channels were dedicated to recording continuous local field potential from V1 in each implanted hemisphere. In a subset of experiments, an additional third recording channel was reserved for the Piezo-electrical input carrying the forepaw movement. Local field potential was recorded from V1 with 1 kHz sampling. On the Plexon system, we used a 500 Hz low-pass filter. On the Intan system, we used a 0.1 Hz high pass and a 7.5 kHz low pass filter. Local field potential data and piezo-electric data was imported (see Importing and data cleaning) and the local field potential’s spectral content was analyzed (see Spectral analysis). In a subset of LFP experiments, forepaw movement was analyzed (see Movement analysis). In a separate LFP experiment, pupil dilation was monitored (see Pupil analysis).

### In vivo two-photon calcium imaging

Three to four weeks following craniotomy surgery, mice were habituated to the behavior restraint apparatus in front of a gray screen with the objective lens of the two-photon microscope positioned on the head plate for 30 min for two consecutive days before beginning their visual stimulus delivery. A Ti:sapphire laser (Coherent, Santa Clara, CA, USA) was used for imaging at a wavelength of 930 nm. Photomultiplier tubes (Hamamatsu, Japan) and the objective lens (20×, 0.95 NA, XLUMPLFLN, Olympus, Japan) were used to detect fluorescence images. Calcium image recordings were triggered by time-locked TTL pulses generated from USB-1208fs (Measurement Computing, Norton, MA, USA) using the Prairie view and TriggerSync Software (Bruker, CA, USA) and imaged at a frequency of ∼2.8 Hz at the depth of ∼350 µm in V1. The size of the imaging field of view was ∼600 x 600 µm^2^ at 256 x 256 pixels.

### Pupillometry

To track the pupil during head fixation, we used a Blackfly S USB3 camera (FLIR Systems, Inc., Wilsonville, OR, US) with a 1.0X lens (Edmund Optics, Inc., Barrington, NJ, US). The left eye was illuminated with a 780 nm infrared LED light source (Thorlabs, Inc., Newton, NJ, US). A small tissue was placed over the light source to disperse luminance. Images were acquired at 20 frames per second during stimulus presentation and each frame emitted a voltage signal into the RHD Recording system for later alignment with stimulus presentations. A subset of videos was used for training the top and bottom edge of the pupil on DeepLabCut (Mathis et al., 2018). All videos were evaluated with the trained network. The output of DeepLabCut includes the x,y-coordinate of the top and bottom edge as well as the certainty of the location. Both were used in our analysis (see Import and data cleaning and Pupil analysis).

### Importing and data cleaning

All analyses were conducted using custom MATLAB code and the Chronux toolbox (Bokil et al., 2010). All code is available upon reasonable request. Briefly, the local field potential from each channel was extracted and converted to microvolts. Data was then zero-meaned and de-trended using a 1 second sliding window and a 500 ms step size. A 3rd-order Butterworth filter was used to notch frequencies between 58 and 62 Hz. The average voltage of the first ten ms after a phase-reversal was subtracted from each individual trace to align them. For average visually-evoked potentials (VEPs), data was smoothed with a gaussian spanning 20 milliseconds using MATLAB’s *smoothdata* function. Piezo-electric data was zero-meaned and rectified. Pupil edges that had less than 100% certainty of location in terms of DeepLabCut output (see Pupillometry) were ignored and a spline interpolation was used to recover the missing points. Qualitatively, these periods of uncertain pupil edge location occurred frequently during black screen and rarely during gray screen or visual stimulus presentation.

### Spectral analysis

Given that the visually-evoked potential violates assumptions required for spectral analysis (namely second-order stationarity), we only analyzed the spectral activity between 400 ms and 2000 ms after a phase-reversal. We computed the multi-tapered spectrogram of the local field potential using the Chronux toolbox (Bokil et al., 2010). The parameters used were: a 500 ms sliding window, a 100 ms step size, zero-padded to the 2nd power, and 5 tapers with a time-bandwidth product of 3. We also computed the multi-tapered spectrum using the same parameters, but including all data between 400 ms and 2000 ms. To calculate the normalized spectrum/spectrogram, we found the median spectrum/spectrogram of the animal’s black screen and took 10*log10(stimulus_spectrum/median_black_spectrum). This is reported as a decibel (dB).

### Concatenated spectrum analysis

Given the contamination by the visually-evoked potential, we concatenated the normalized spectrums. This concatenated spectrum utilizes the multi-tapered spectrum of the period between 400 and 2000 ms after a phase-reversal. Ordering these spectrums by their presentation number and representing power as a color generated the concatenated spectrum. For each presentation, we calculated the max power within the 10-30 Hz frequency band as well as the 60-80 Hz frequency band. By visual inspection, no changes were seen in the concatenated spectrum after 25 presentations, so we used the average of the max power for presentations 26-100 as a metric to compare the first few presentations in our bootstrapping procedure. No significant difference is found between presentations after the 15^th^ and the average of presentations 26-100, confirming that this data split was reasonable. Other splits were tried and there were no qualitative differences in the resulting data.

### Block onset spectral analysis

For this analysis, we only used the group of animals that had 6 second gray periods and 10 second black periods (see Stimulus delivery). The local field potential data within 12 seconds of block onset (both before and after) was extracted and separated into overlapping 400 ms chunks each with centers spaced 100 ms apart. For each chunk, we computed the normalized spectrum (see Spectral analysis). Transitions from black to gray, gray to stimulus, and between phase 0° and phase 180° will elicit a visually-evoked potential. Thus, for each frequency within the normalized spectrum, we removed contaminated regions and interpolated between them using cubic interpolation. Specifically, we removed the chunks whose midpoints were between 100 ms before and 500 ms after a transition. The trailing edge (near 12 seconds) was not included due to the edge artifacts of interpolation. Once we had the interpolated normalized spectrum, we found the max power in the 10-30 Hz frequency band and the 60-80 Hz frequency band for each chunk.

### P-Episode analysis

P-Episode is a method that quantifies the fraction of time that oscillations exceed amplitude and duration thresholds (Caplan et al., 2001; van Vugt et al., 2007). We lightly adapted analysis software kindly provided by Marieke van Vugt (Univeristy of Groningen). Briefly, Mortlet wavelets between 7 and 100 Hz with a wavenumber of 5 were used to extract spectral information from both black and stimulus periods. Then, to obtain an estimate of the background spectral activity, we fit the black spectrum with a linear regression in log-log space and stored the mean power values at each frequency (using the trained regression parameters). This model has the form A/f^α^ and is commonly called “pink” or “colored” noise (Caplan et al., 2001; van Vugt et al., 2007). This was done for each black period and ultimately averaged to get one estimate of the background spectrum per animal. In line with default parameters, the power threshold was determined for each frequency as the 95th percentile of the chi-squared probability distribution with 2 degrees of freedom. The duration threshold was simply three cycles. Next, for each presentation of a stimulus, the above Mortlet wavelets were used to extract spectral information. Finally, for each time point, it was determined whether a given frequency exceeded both the power and duration threshold. If it exceeded both thresholds, the time point was in that oscillation. Otherwise, the time point was not in that oscillation. Herein we report the percent of time in an oscillation for each of the analyzed frequencies.

### Correlations

We analyzed the correlation of VEP magnitude and LFP power in single trials. Because the VEP in response to each phase reversal is variable and can be obscured by ongoing voltage fluctuations, it had to exceed a threshold to be included in the analysis. For each animal, we computed the average activity during exposure to gray screen between each block of stimuli. The gray period was sampled at the same frequency and duration as used for VEP analysis. The difference between the minimum of this average gray activity (within 100 ms of the sample onset) and the maximum of this average gray activity (any time after the minimum) was taken as our voltage threshold. Next, we computed the average VEP above threshold for all familiar and novel presentations to get the indices for the positive and negative peaks (regardless of stimulus). Using these indices, for each phase-reversal, we calculated the magnitude of the difference between the positive and negative peak. If this single-trial VEP magnitude was below the voltage threshold decided by the gray screen period, that trial was discarded (∼19% of trials). We additionally eliminated the first presentation of each block since we were comparing the pre-phase-reversal LFP to the VEP magnitude and the first presentation’s LFP would be during a gray screen.

For each phase-reversal, we also calculated the normalized spectrum for the 400 ms leading into the phase-reversal. Using this, we obtained the max power within the 10-30 Hz frequency band and the 60-80 Hz frequency band. For correlation analysis, we used a simple Pearson correlation coefficient. Correlation analysis could be done on just familiar data, just novel data, or an equal random sampling of both.

### Calcium imaging analysis

Acquired time series of calcium imaging files were processed using Suite2p (Pachitariu et al., 2017). All recorded files were registered to stabilize shifts due to animal movement. We manually selected regions of interest (ROIs) base on the maximum projection of all frames. We then extracted the fluorescence of each ROI for all time points. In line with previous work, for each ROI we calculated the estimated true fluorescence of the ROI. This is the measured fluorescence of the ROI minus 7/10ths of the average measured fluorescence of the surrounding neuropil (Chen et al., 2013). We used the average inter-block gray period to compute the response relative to gray (F_stim_ – F_avg_gray_)/F_avg_gray_. For our non-parametric bootstrapping procedure, we randomly selected with replacement from the mice we recorded from, randomly selected with replacement from the cells they had, and randomly selected with replacement from said cell’s data. This procedure was repeated as outlined in **Statistics**. Only cells that could be tracked over all days were included in our analysis.

### Movement analysis

For each stimulus phase-reversal, we smoothed imported piezo-electric data with a moving average over 200 ms. We similarly smoothed periods of gray screen (see Stimulus delivery). We subtracted the median gray screen forepaw movement from the stimulus forepaw movement and report this as normalized movement in arbitrary units (a.u.).

### Pupil analysis

The DeepLabCut predicted x,y-coordinates of the top and bottom of the pupil were cleaned (see Importing and data cleaning). Then, the Euclidian distance between the top and bottom of the pupil was calculated to obtain the pupil dilation in pixels.

### Statistics

Most statistics were done with the non-parametric hierarchical bootstrap for multi-level data (Saravanan et al., 2020). Briefly, statistical comparisons were between two groups (with each animal belonging to both groups due to the within-animal experimental design). To begin the bootstrap process, mice were randomly selected with replacement from the population. For each randomly selected mouse, a number of random trials were selected with replacement from the mouse’s group A data. Another random set of trials were selected with replacement from the mouse’s group B data. This data was stored and the process was repeated for each randomly selected animal. In some instances (**Figure 4I-J, Figure 7, Figure 8, Figure 9**, and **Figure 12**), blocks were randomly selected with replacement and all trials or timepoints within that block were used. Once all data was randomly selected, the mean difference between the randomly selected samples of group A and group B was computed and stored. This entire bootstrap process was repeated 1000 times. Once all 1000 bootstraps had been completed, the bootstrapped differences were sorted from lowest to highest value. The 500th value was the median group difference, the 5th value was the lower bound of the 99% confidence interval, and the 995th value was the upper bound of the 99% confidence interval. If the 99% confidence interval does not include zero, we report a statistically significant difference between group A and group B with a small marker below the corresponding data on the plot. To facilitate communication in the results section, we report the identity, median, and 99% confidence interval for the peak significant median difference above and below 50 Hz, if it exists. In most figures, the colored plots are mean +/-SEM of all animals and the gray plots to the right are the 99% bootstrapped confidence interval for those two groups. The only other statistical procedure was a two-sample Kolmogorov-Smirnov test on the data comprising the cumulative distribution functions in **Figure 5**. All data, code, and values are available for complete replication of the paper upon reasonable request.

**Figure 1.**
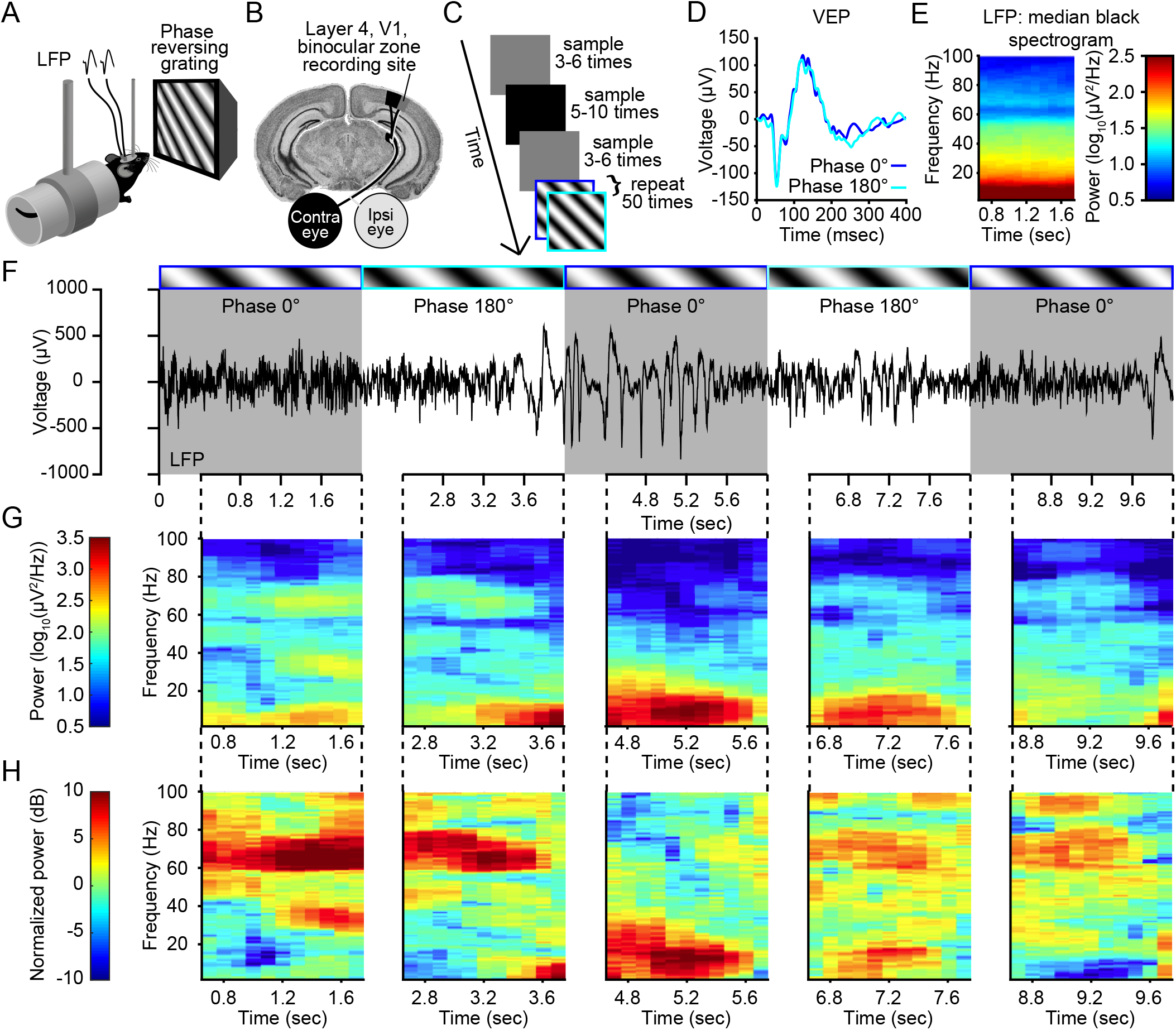
Layer 4 local field potential displays variable frequency composition in V1 of awake, head-fixed mice: (**A**) We recorded local field potentials (LFP) from primary visual cortex (V1) in awake, head-fixed mice in response to phase-reversing sinusoidal grating stimuli. (**B**) Electrodes were chronically implanted bilaterally in thalamo-recipient layer 4 of binocular V1. (**C**) The experimental set-up (see Methods). (**D**) The average VEPs for the phase 0° (“flip”, blue trace) and phase 180° (“flop”, cyan trace) stimuli recorded in V1 of the example mouse for which LFPs are presented in panels **E-G**. (**E**) The median spectrogram for a period of black screen activity corresponding to similar periods for phase-reversing grating stimuli displays the expected inverse power-frequency relationship (a.k.a. “pink noise”) common in neural recordings, but otherwise reveals no time-dependent dynamics in spectral power. (**F**) Examination of the continuous LFP (black trace) relative to each phase-reversal (gray and white bars) reveals marked variability in V1 activity. Blue and cyan colored outlines around a visual stimulus identify the grating phase as either 0° (“flip”) or 180° (“flop”). The dashed black line connecting the time series trace with the spectrogram indicates the period of time spectral analysis was conducted on. (**G**) The raw spectrogram for the periods outlined in (**F**). (**H**) The normalized spectrogram for the same periods outlined in (**F**).

**Figure 2.**
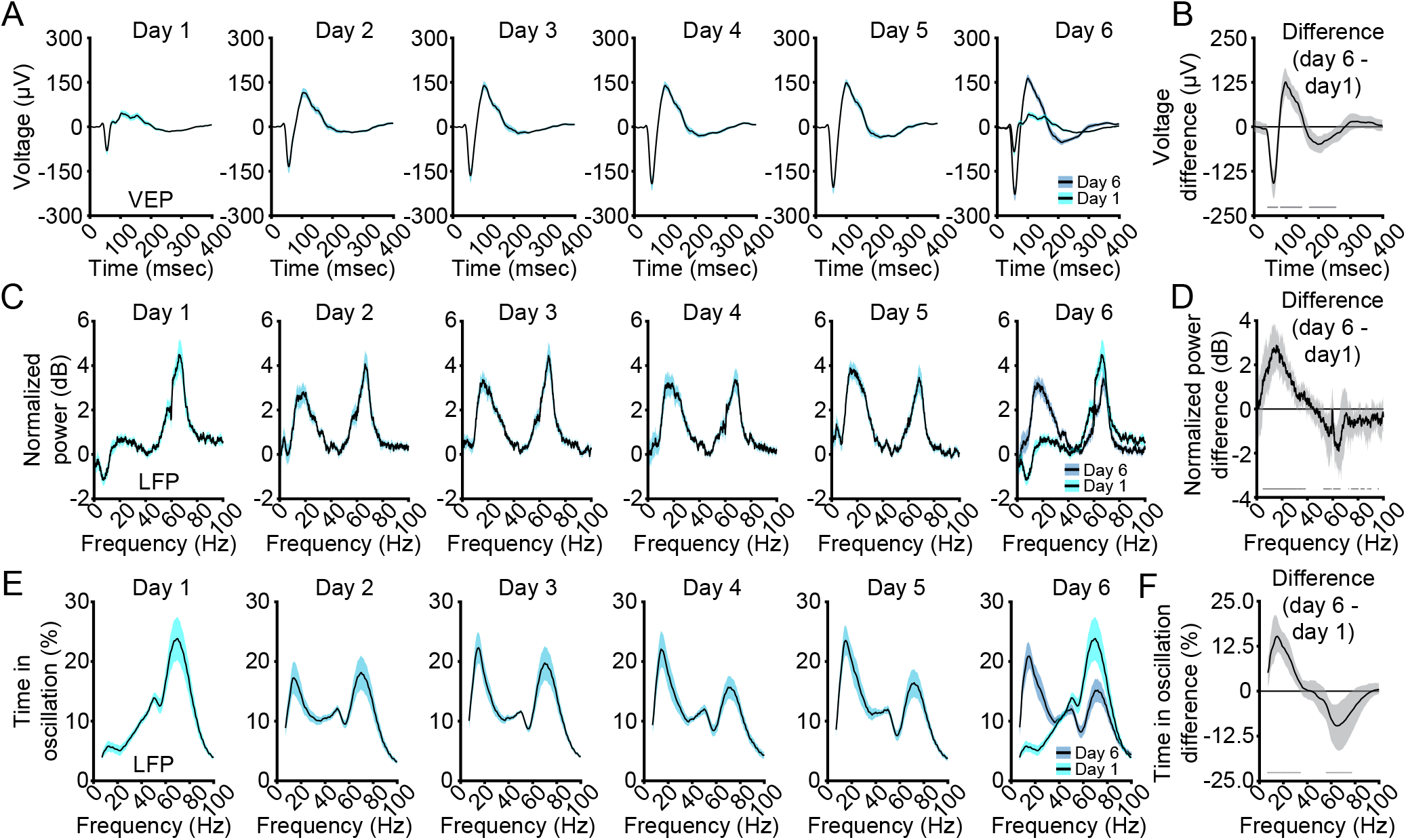
V1 VEPs and LFP oscillations are influenced by stimulus familiarity over days: (**A**) Repeated presentation of the same stimulus orientation over 6 days increases the average VEP (n = 13). (**B**) Non-parametric hierarchical bootstrapping results confirm the VEP on day 1 (cyan) is significantly less than day 6 (blue), despite the fact the stimulus has not changed. Solid line represents the median value and the shaded region reflects the 99% confidence interval. (**C**) Over those same 6 days in the same mice, low frequency power increases and high frequency power decreases. (**D**) Non-parametric hierarchical bootstrapping results confirm that the spectrum on day 6 (blue) is significantly different from the spectrum on day 1 (cyan), despite the fact the stimulus has not changed. Solid line represents the median value and the shaded region reflects the 99% confidence interval. (**E**) Analysis of the amount of time the LFP shows sustained oscillatory activity for a given band. (**F**) Non-parametric hierarchical bootstrapping results of time differences. Marks near the x-axis in (**B**), (**D**), and (**F**) indicate the 99% confidence interval does not include 0 (thus the difference is statistically significant). Averaged VEPs, spectrums, and P-Episode results are presented as mean +/-SEM.

**Figure 3.**
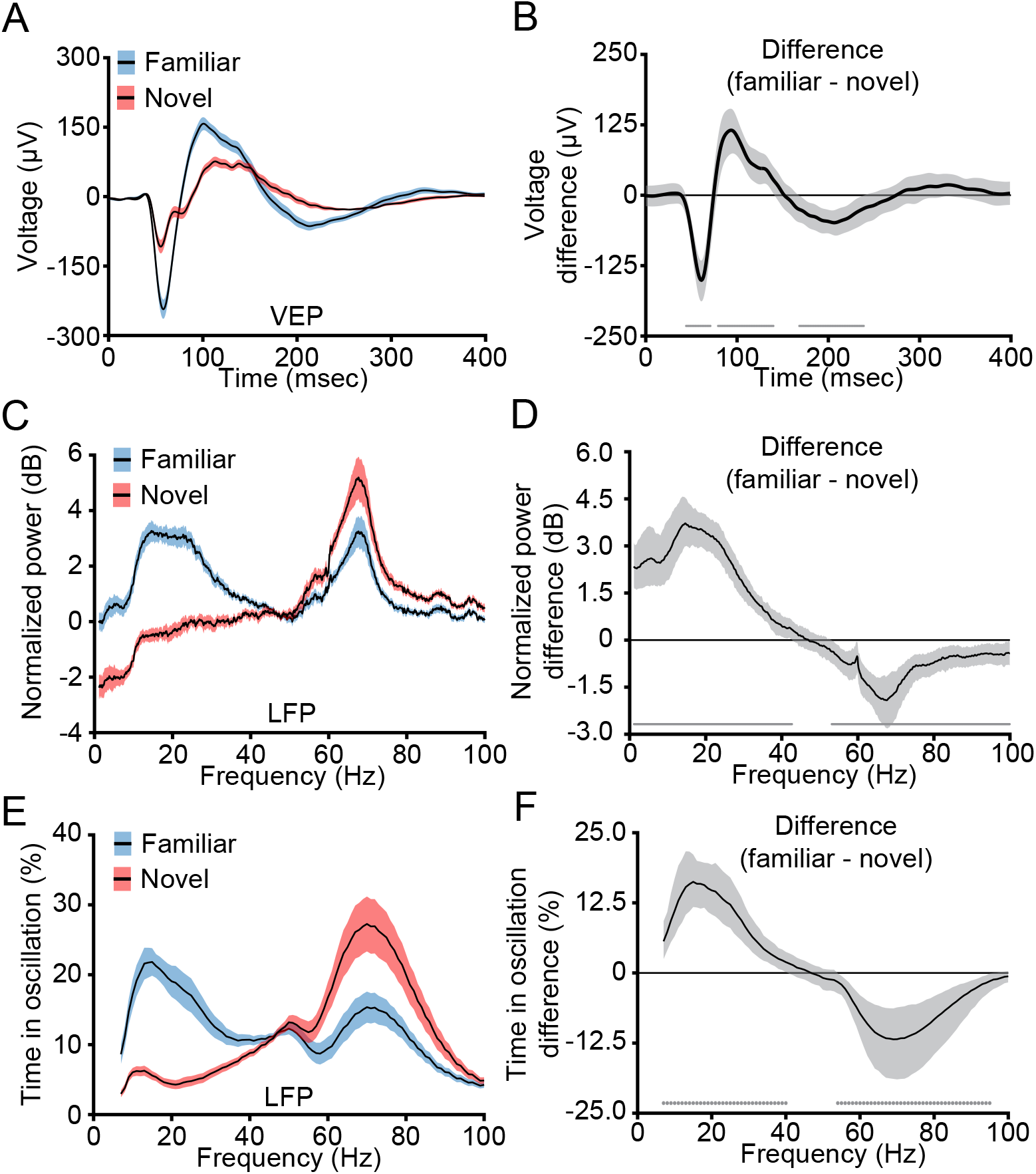
Experience-dependent changes in V1 VEPs and LFP oscillations are stimulus-specific: (**A**) Presentation of a novel (red) stimulus elicits a smaller visually-evoked potential (VEP) than a familiar (blue) stimulus (n = 13). (**B**) Non-parametric hierarchical bootstrapping results confirm that the familiar VEP is significantly larger than the novel VEP. Solid line represents the median value and the shaded region reflects the 99% confidence interval. (**C**) In the same mice, presentation of a novel (red) stimulus increases high frequency power and decreases low frequency power compared to a familiar (blue) stimulus. (**D**) Non-parametric hierarchical bootstrapping results confirm that the familiar spectrum is significantly different from the novel spectrum. Solid line represents the median value and the shaded region reflects the 99% confidence interval. (**E**) Analysis of the time the LFP shows sustained oscillatory activity for a given band. (**F**) Non-parametric hierarchical bootstrapping results. Marks near the x-axis in (**B**), (**D**), and (**F**) indicate the 99% confidence interval does not include 0 (thus the difference is statistically significant). Averaged VEPs, spectrums, and P-Episode results are presented as mean ± SEM.

**Figure 4.**
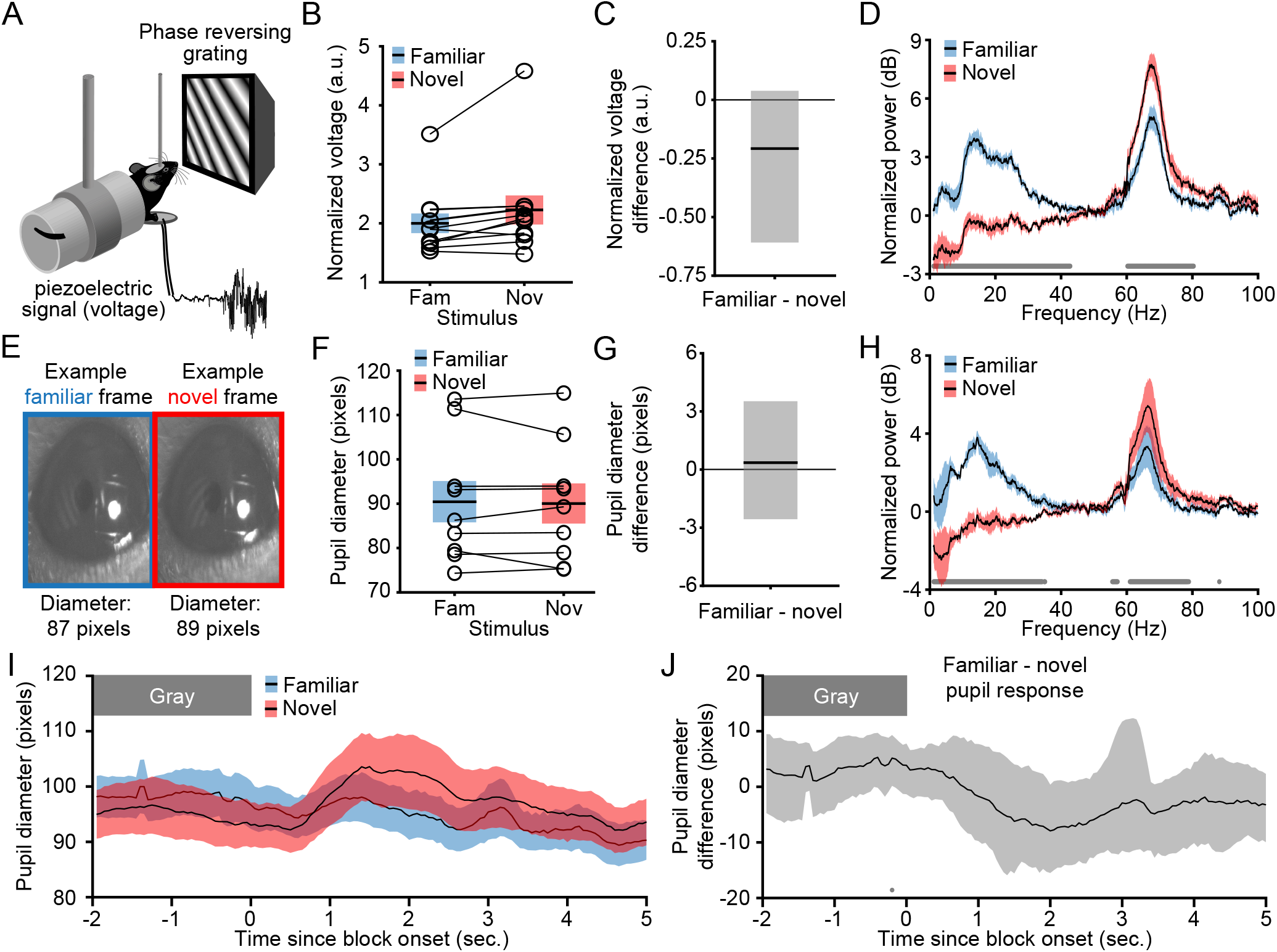
Movement and arousal do not account for the changes in layer 4 LFP frequency composition: (**A**) We recorded forepaw movements of awake, head-fixed mice in response to phase-reversing sinusoidal grating stimuli. (**B**) The normalized piezo-electric voltage indicates there is no difference in the average forepaw movement during blocks of familiar (blue) and novel (red) stimuli over the time when LFPs are analyzed (n = 11). Averaged voltages are presented as mean ± SEM. Open circles with connected lines show average normalized voltage from each animal. (**C**) Non-parametric hierarchical bootstrapping results confirm that there is no piezo-electric voltage difference between familiar and novel stimuli. (**D**) In a subset (5) of these 11 mice, we simultaneously recorded the movement and LFP data. Presentation of a novel (red) stimulus increased high frequency power and decreased low frequency power compared to a familiar (blue) stimulus. Non-parametric hierarchical bootstrapping results confirm that the familiar spectrum is significantly different from the novel spectrum (*marks near the x-axis indicate where the 99% confidence interval does not include 0*). Averaged spectrums are presented as mean ± SEM. (**E**) Pupil size was measured as mice viewed blocks of familiar and novel stimuli. The familiar and novel frame from this exemplar mouse show the approximate population average size and location of the pupil. (**F**) The pupil diameter is similar as mice view familiar (blue) and novel (red) blocks of stimuli over the time when LFPs are analyzed (n = 9). Averaged pupil diameters are presented as mean ± SEM. Open circles with connected lines show individual animal’s average pupil diameter. (**G**) Non-parametric hierarchical bootstrapping results confirm that there is no pupil diameter difference between familiar and novel stimuli. (**H**) Same analysis of LFPs as in (**D**), but with a subset (4) of these 9 mice in which we simultaneously recorded the pupillometry data and LFP data. (**I**) Pupil diameter at block onset (n = 9). Averaged pupil diameters are presented as mean ± SEM. (**J**) Non-parametric hierarchical bootstrapping results show that the familiar pupil diameter and novel pupil diameter are similar. Solid line represents the median value and the shaded region reflects the 99% confidence interval. Marks near the x-axis indicate the 99% confidence interval does not include 0 (thus the difference is statistically significant). The one statistically significant point in (**J**) is before the stimulus train appears and is likely a Type I error (false positive).

**Figure 5.**
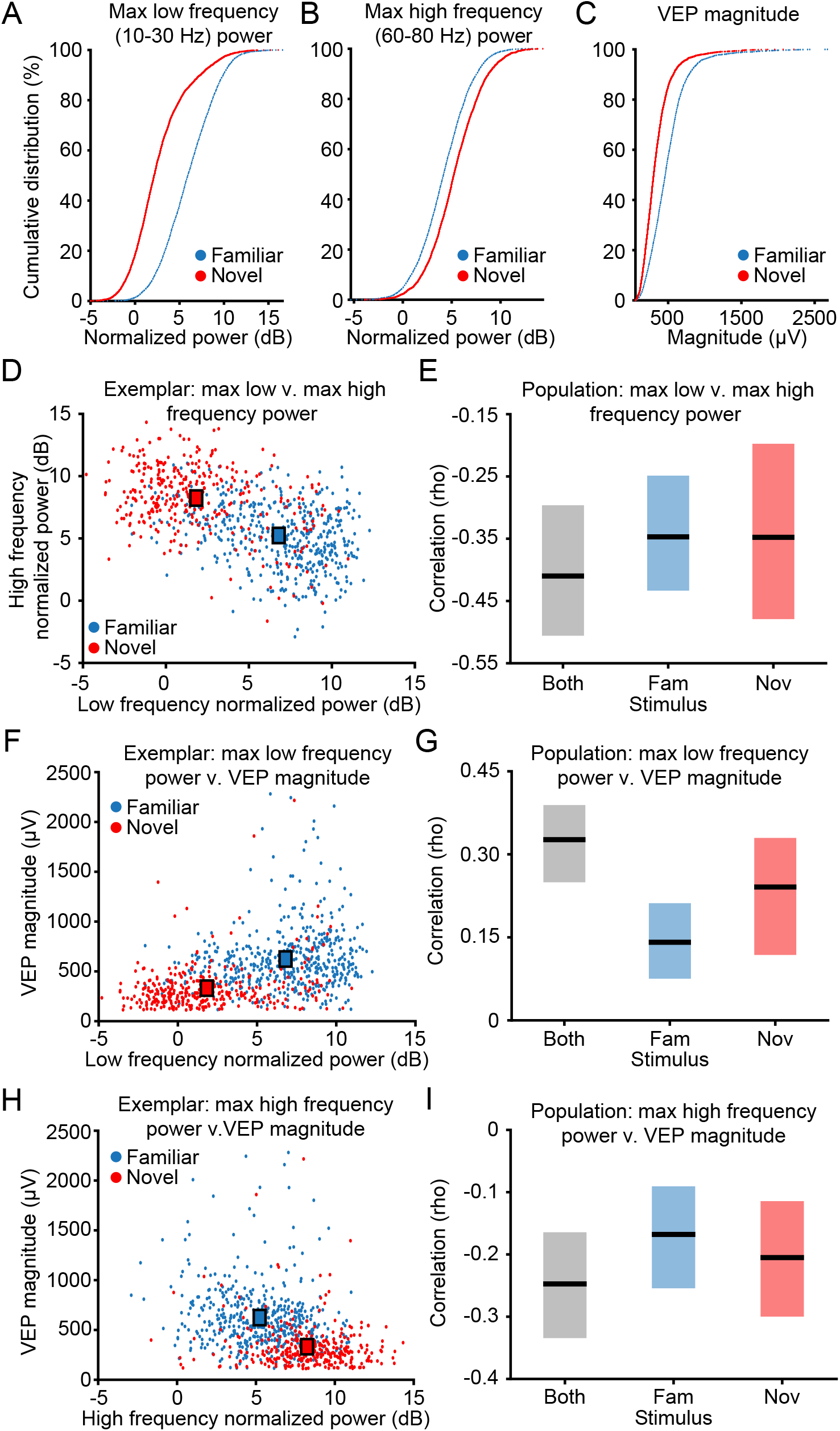
Oscillations and VEP magnitude correlate: (**A**) LFPs and VEPs proximal to 500 stimulus phase reversals were studied in 13 mice. The max low frequency (10-30 Hz) power is larger for familiar stimuli than novel stimuli. (**B**) The max high frequency (60-80 Hz) power is larger for novel stimuli than familiar stimuli. (**C**) The VEP magnitude is larger for familiar stimuli than novel stimuli. (**D**) An exemplar animal’s scatterplot of each presentation’s max low frequency power and high frequency power shows they are negatively correlated. (**E**) Non- parametric hierarchical bootstrapping results confirm that low frequency and high frequency power are negatively correlated, regardless of stimulus. (**F**) Same as in (**D**), but for low frequency power and VEP magnitude. (**G**) Same as in (**E**), but showing low frequency power positively correlates with VEP magnitude. (**H**) Same as in (**D**), but for high frequency power and VEP magnitude. (**I**) Same as in (**E**), but showing high frequency power negatively correlates with VEP magnitude. Solid lines in (**E**), (**G**), and (**I**) indicate the median value and the shaded region reflects the 99% confidence interval. “Both” refers to a correlation analysis done with equal random sampling of both familiar and novel stimulus presentations. Squares in (**D**), (**F**), and (**H**) represent the midpoint of each group’s scatterplot cloud.

**Figure 6.**
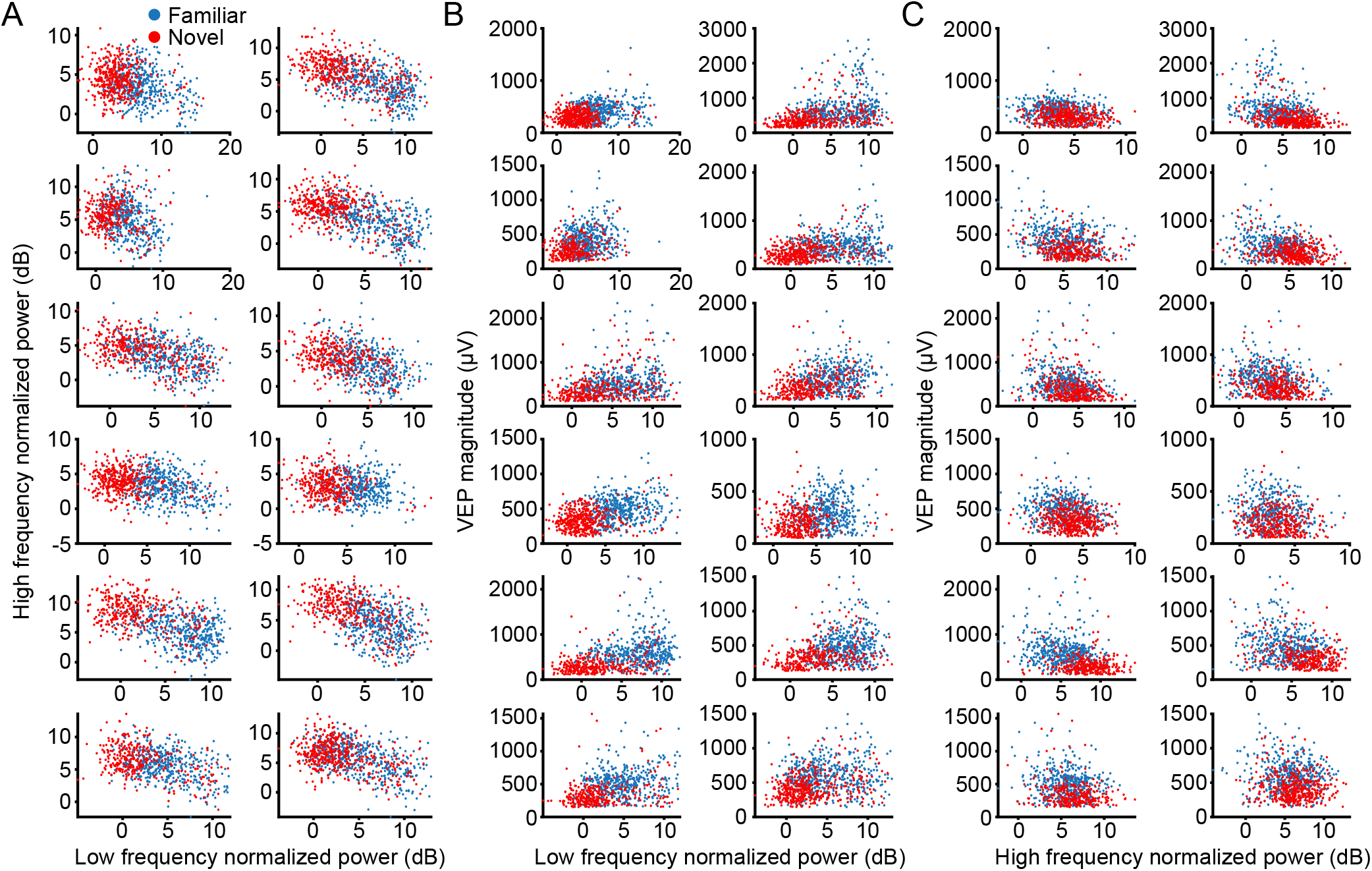
Animal breakdown of correlation of VEP and oscillations: (**A**) Scatterplots of max low frequency (10-30 Hz) power and max high frequency (60-80 Hz) power for 12 of the 13 mice (the 13^th^ is the exemplar in Figure 5D). (**B**) Scatterplots of max low frequency power and VEP magnitude for 12 of the 13 mice (the 13^th^ is the exemplar in Figure 5F). (**C**) Scatterplots of max high frequency power and VEP magnitude for 12 of the 13 mice (the 13^th^ is the exemplar in Figure 5H).

**Figure 7.**
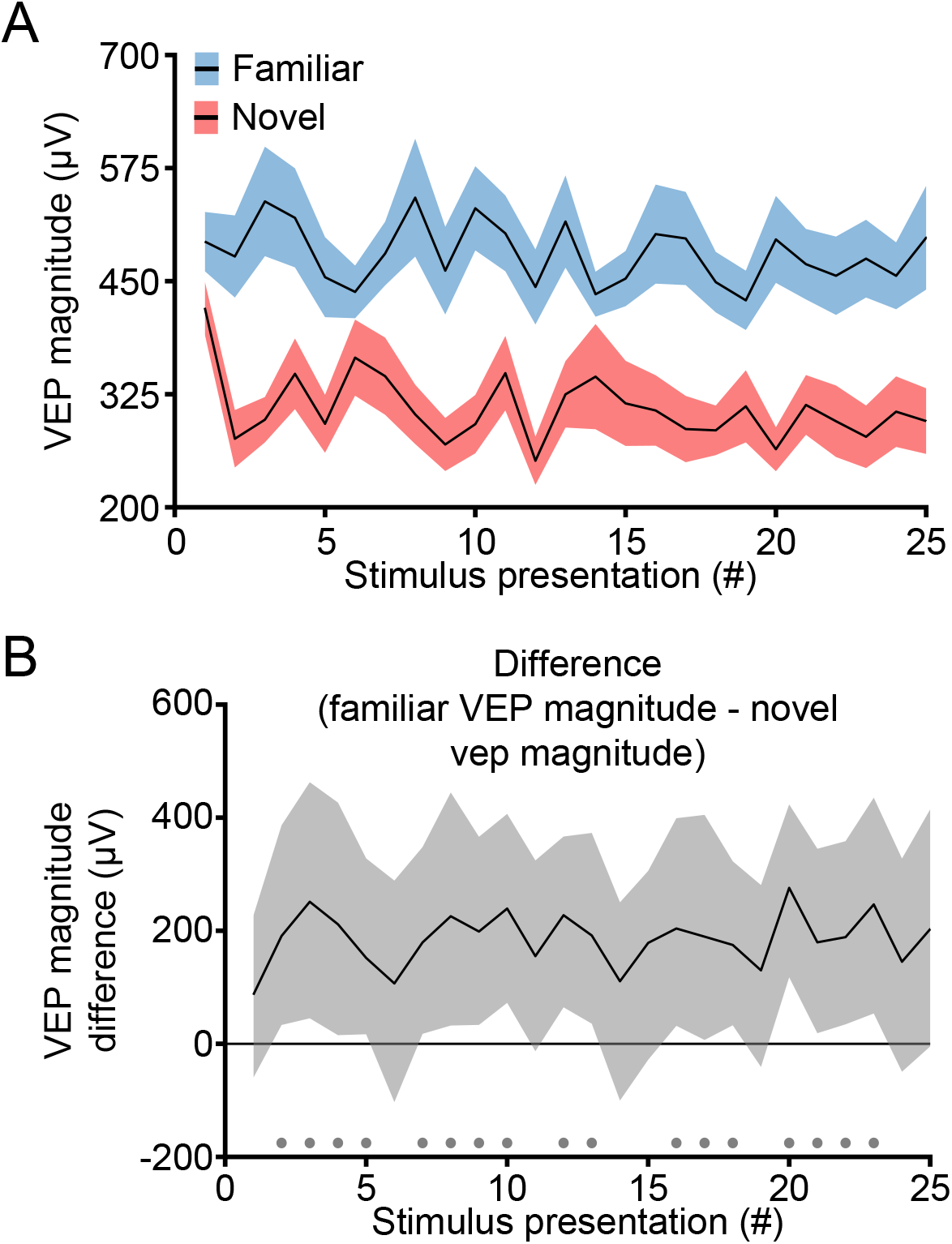
Differences in VEPs to familiar and novel stimuli emerge during a stimulus block: (**A**) In the 13 mice used for LFP analysis, we plot the change in VEP magnitude elicited by familiar and novel stimulus over the early trials from block onset. Presentation 1 corresponds to the VEP elicited by the transition from a gray screen to an oriented grating stimulus, and subsequent presentations correspond to phase-reversals of this grating. Averaged VEP magnitude is presented as mean ± SEM. (**B**) Non-parametric hierarchical bootstrapping results confirm previous findings (Kim et al., 2019) that VEP differences to familiar and novel stimuli only emerge after presentation 1. Marks near the x-axis indicate trials with a statistically significant difference.

**Figure 8.**
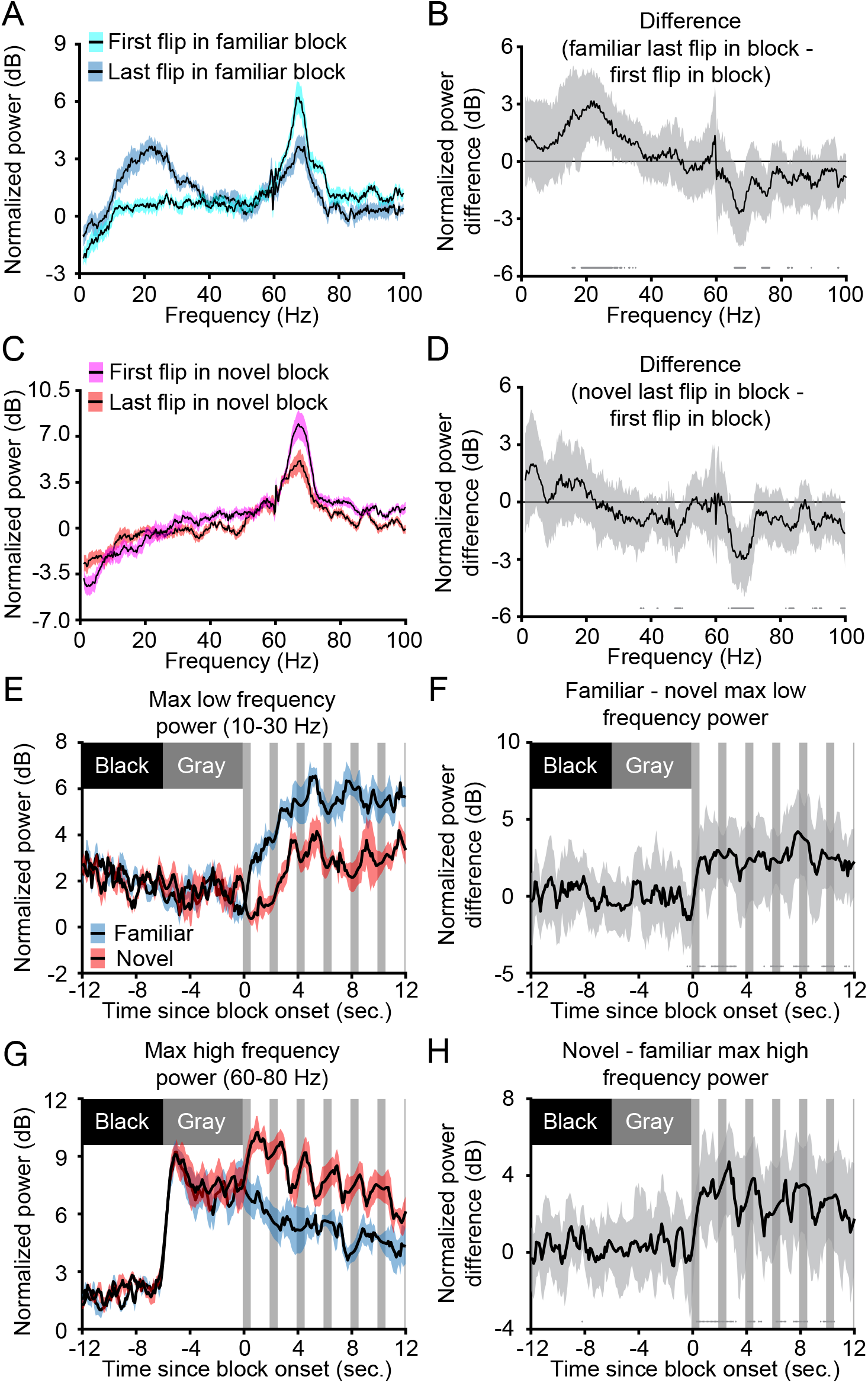
Experience-dependent oscillations in V1 change within a stimulus block: (**A**) The LFP following the onset of a familiar stimulus block (cyan) has increased high frequency power and decreased low frequency power compared to the last presentation of the same phase stimulus (blue). Averaged spectrums (n = 13) are presented as mean ± SEM. (**B**) Non-parametric hierarchical bootstrapping results confirm that the last flip’s spectrum is significantly different from the first flip’s spectrum. Solid line represents the median value and the shaded region reflects the 99% confidence interval. (**C**) The onset of a novel stimulus block (magenta) has increased high frequency power compared to the last contrast reversal of the same phase (red). (**D**) Non-parametric hierarchical bootstrapping results for (**C**). (**E**) The max low frequency power near block onset increases to a higher value for familiar stimuli compared to novel stimuli. Gray vertical bars represent periods where the LFP is contaminated by the VEP so interpolation was done to connect non-contaminated regions of the plot (see Methods). Averaged power is presented as mean ± SEM. (**F**) Non-parametric hierarchical bootstrapping results confirm that familiar stimuli have more low frequency power than novel. (**G**) Max high frequency power near block onset increases to a larger value for a novel stimulus than a familiar one. (**H**) Same as in (**F**), but focused on high frequency power at block onset. Marks near the x-axis in (**B**), (**D**), (**F**), and (**H**) indicate the 99% confidence interval does not include 0 (thus the difference is statistically significant).

**Figure 9.**
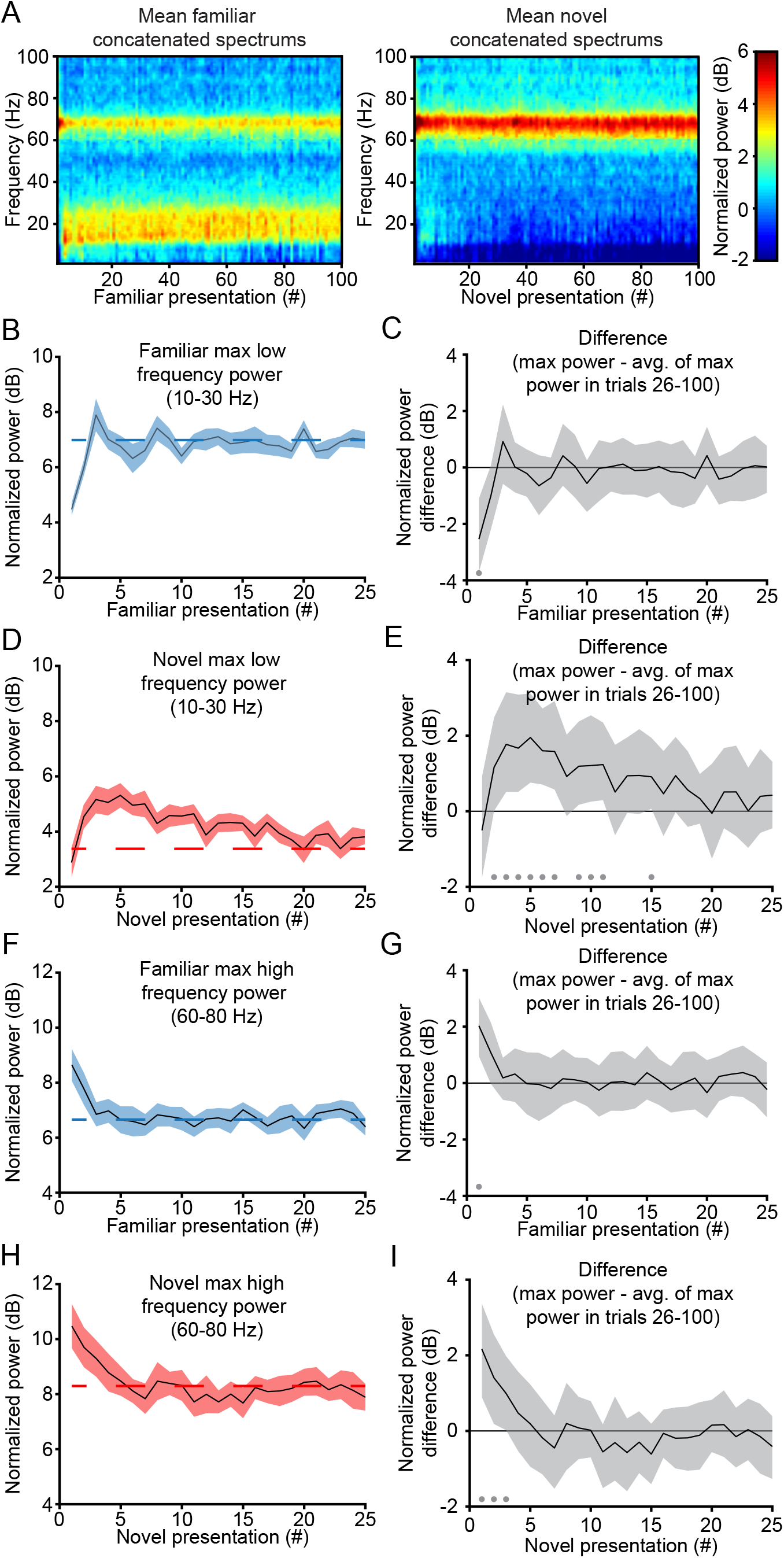
Emergence of experience-dependent oscillations in V1 during blocks of stimulation: (**A**) The average concatenated spectrum (see Methods) reveals short-term dynamics in oscillatory power for both familiar and novel stimuli (n = 13). (**B**) For familiar stimuli, the max power in the 10-30 Hz band quickly increases to a steady value after the first presentation. (**C**) Non-parametric hierarchical bootstrapping results confirm that only the first presentation shows significantly different changes from the average of the max power in the last 75 phase-reversals. (**D**) For novel stimuli, the max power in the 10-30 Hz band rises during the 2^nd^ through 5^th^ presentations and slowly decays over the next 5-10 presentations. (**E**) Non-parametric hierarchical bootstrapping results confirm the significantly different spectral changes from the average of the max power in the last 75 phase-reversals outline in (**D**). (**F-G**) Same as (**B-C**), but showing, for familiar stimuli, that the max power in the 60-80 Hz band quickly decreases to a steady state value after the first presentation. (**H-I**) Same as (**D-E**), but showing, for novel stimuli, that the max power in the 60-80 Hz band decreases to a steady state value after the first few presentations. Spectral power in (**B**), (**D**), (**F**), and (**H**) are presented as mean +/-SEM and the dashed red line represents the average of the max power in the last 75 phase-reversals. Marks near the x-axis in (**C**), (**E**), (**G**), and (**I**) indicate the 99% confidence interval does not include 0 (thus the difference is statistically significant). Additionally, the solid line in (**C**), (**E**), (**G**), and (**I**) represents the median value and the shaded region reflects the 99% confidence interval.

**Figure 10.**
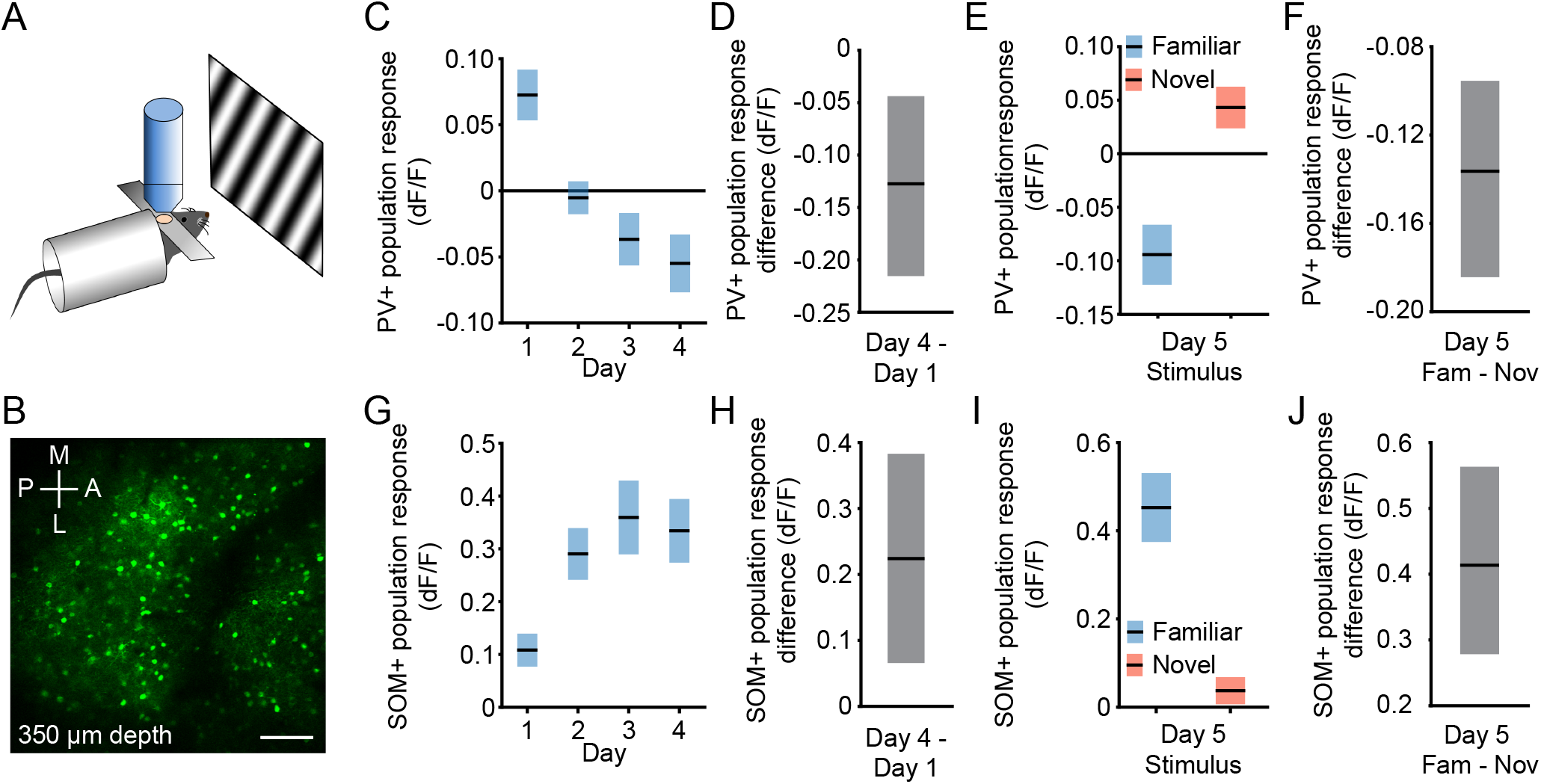
PV+ and SOM+ neurons in layer 4 respond differently to blocks of familiar and novel stimuli: (**A**) We recorded parvalbumin-expressing (PV+) cell activity or somatostatin-expressing (SOM+) cell activity from layer 4 of V1 in awake, head-fixed mice in response to phase-reversing sinusoidal grating stimuli. (**B**) Image shows an example field of view (scale bar, 100 µm). (**C**) PV+ cell activity gradually diminishes with stimulus experience (mean ± SEM, n = 9). (**D**) Non-parametric hierarchical bootstrapping results confirm day 4 had less activity than day 1. (**E**) On day 5, PV+ cell activity is elevated while the mouse views novel stimuli. (**F**) Non-parametric hierarchical bootstrapping results confirm familiar stimuli induce less PV+ cell activity than novel stimuli. (**G**) SOM+ cell activity increases with stimulus experience (mean ± SEM, n = 7). (**H**) Non-parametric hierarchical bootstrapping results confirm day 4 had more activity than day 1. (**I**) On day 5, SOM+ cell activity is elevated while the mouse views familiar stimuli. (**J**) Non-parametric hierarchical bootstrapping results confirm familiar stimuli induce more SOM+ cell activity than novel stimuli. All data is reported relative to the average inter-block gray screen activity (see Methods). Activity is averaged across all cells for each animal and presented as the group mean +/-SEM in (**C**), (**E**), (**G**), and (**I**). Solid black lines in (**D**), (**F**), (**H**), and (**J**) indicate the median value for the difference and the shaded regions reflect the 99% confidence interval.

**Figure 11.**
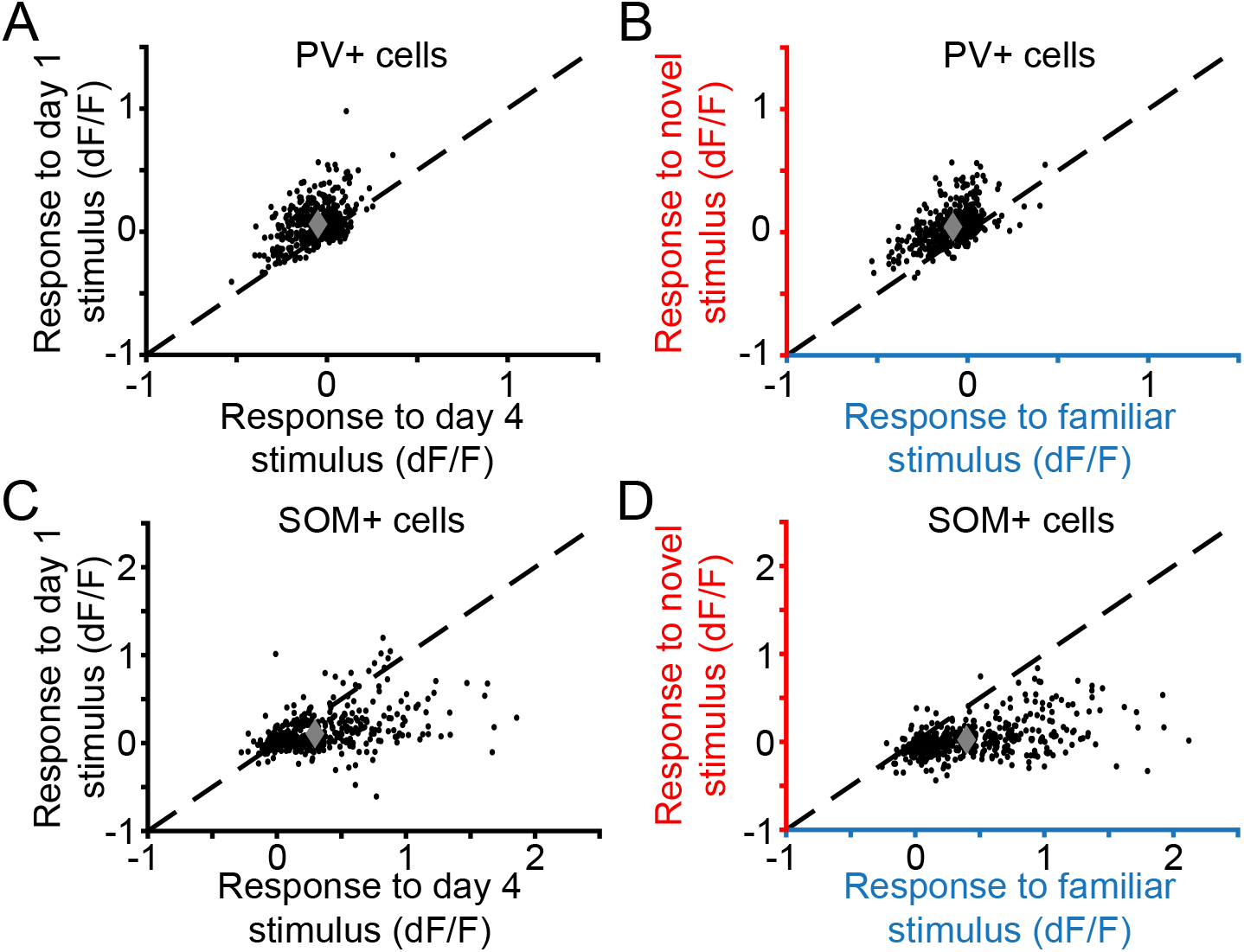
Familiar-novel differences in activity of parvalbumin-expressing and somatostatin-expressing neurons are remarkably uniform: (**A**) Plotted are the responses of each PV+ neuron recorded during visual stimulation with a phase reversing grating of the same orientation on the first and 4^th^ day (n = 1,251 PV+ neurons from 9 mice) . (**B**) Responses of the same PV+ neurons on day 5, comparing the familiar and novel stimulus orientations. (**C**) Responses of each SOM+ cell plotted on day 1 vs day 4 of viewing the same oriented stimulus (n = 1,021 SOM+ neurons from 7 mice). (**D**) Responses of the same SOM+ neurons on day 5, comparing the familiar and novel stimulus orientations. All data reported relative to the average inter-block gray screen activity (see Methods). Dashed line in (**A**-**D**) is the identity line y = x.

**Figure 12.**
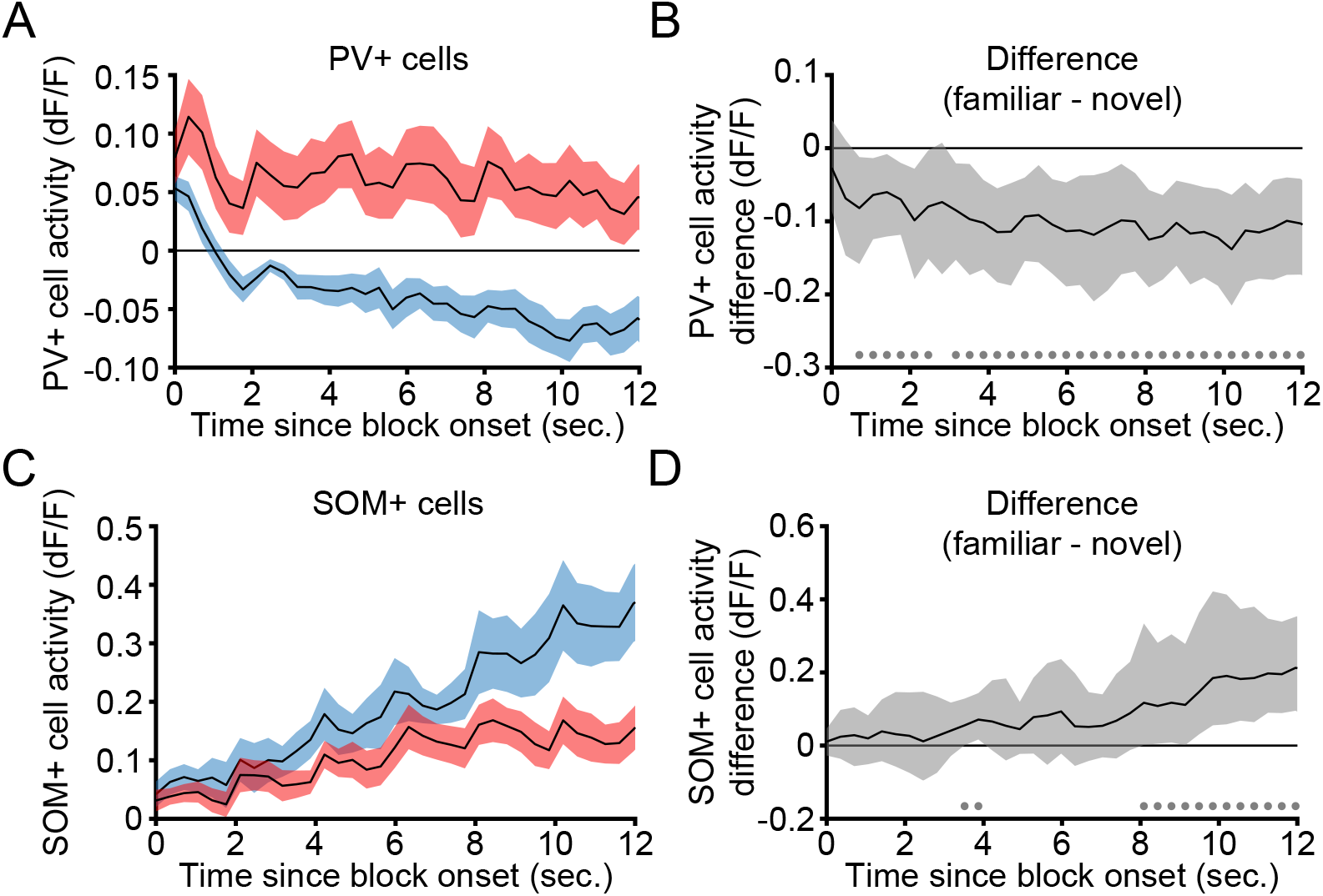
Experience-dependent differences in the activity of PV+ and SOM+ emerge during blocks of stimulation: (**A**) PV+ cell activity from layer 4 of V1 in 9 awake, head-fixed mice in response to the transition from gray screen (time 0) and during subsequent phase-reversals of sinusoidal grating stimuli every 2 sec. Both familiar (blue) and novel (red) showed an increase in PV+ cell activity at block onset followed by a decrease that was more pronounced during familiar stimulus viewing. (**B**) Non-parametric hierarchical bootstrapping results confirm that PV+ cell activity during novel blocks is larger than familiar blocks as early as 1 second after stimulus onset. (**C-D**) Same as in (**A-B**) but measuring SOM+ cell activity in 7 mice. SOM+ cell activity is increased after block onset during stimulation with both familiar (blue) and novel (red) gratings. However, the increase during novel stimulus viewing is less than during familiar stimulus viewing. Non-parametric hierarchical bootstrapping results confirm that SOM+ cell activity during familiar blocks is larger than novel blocks as early as 3-4 seconds. Activity is averaged across all cells for each animal and presented as the group mean ± SEM in (**A**) and (**C**). Solid black lines in (**B**) and (**D**) indicate the median value for the difference and the shaded regions reflect the 99% confidence interval. Marks near the x-axis in (**B**) and (**D**) indicate the 99% confidence interval does not include 0 (thus the difference is statistically significant).

## Results

### Layer 4 local field potential oscillations display variable frequency composition in V1 of awake, head-fixed mice

We acquired local field potential (LFP) data from electrodes chronically implanted within layer 4 of binocular V1 of C57BL/6 mice. Awake, head-fixed mice viewed full field, 0.5 Hz phase-reversing sinusoidal grating stimuli separated into blocks of 100 phase-reversals preceded by periods of gray and black screen (**Figure 1A-C**). We used equally-spaced timestamps to segment gray and black screen data into 2000 ms portions for further normalization and comparison. Under these conditions, we could average the stimulus-evoked LFP waveform occurring within a 400-ms time window from the start of each phase-reversal (**Figure 1D** shows this for a representative mouse). This average VEP is the signal typically used to monitor the emergence of SRP (Frenkel et al., 2006; Cooke and Bear, 2010; Cooke et al., 2015; Kaplan et al., 2016). However, the continuous LFP signal reveals periodic changes from low amplitude, high frequency activity to high amplitude, low frequency activity (**Figure 1F** shows this for a representative mouse). Because the portion of the recording containing the VEP violates second-order stationarity, a requirement for oscillatory analysis, we focused our analysis on the last 1600 ms of each 2000 ms presentation (**Figure 1F**). This approach is consistent with previous work (Chalk et al., 2010; Zhou et al., 2016). The raw spectrogram for each phase-reversal (**Figure 1F**) shows that some time periods have strong low frequency oscillations whereas others have strong high frequency oscillations (**Figure 1G**). We chose not to normalize the raw signal to data from a gray screen period because isoluminant gray screens elicit narrow-band oscillations at 60 Hz that emerge in the cortex but arise from a subcortical source (Saleem et al., 2017). Instead, we normalized the raw spectrogram to the median spectrogram generated during black screen presentation (**Figure 1E**). The normalized spectrogram for each phase-reversal (**Figure 1F**) again shows periods of strong low frequency oscillations in the alpha/beta range (10-30 Hz) and periods of high frequency oscillations in 60-80 Hz range (**Figure 1H**). As there is unfortunate inconsistency in how the term “gamma” is used in the literature to describe oscillations in visual cortex (Chen et al., 2017; Veit et al., 2017), we have avoided use of this term to describe our findings. However, we note that activity in the 60-80 Hz range is conventionally defined as “high-gamma”. The normalized spectral data will be used throughout the remainder of this study.

### V1 oscillations are influenced by stimulus familiarity over days

We investigated whether the frequency composition of the V1 LFP in layer 4 changes as a result of visual experience. We induced SRP by exposing mice to a phase-reversing stimulus at a single orientation each day for 6 consecutive days. As reported in previous studies (see, e.g., (Frenkel et al., 2006; Cooke et al., 2015; Fong et al., 2020), the VEP magnitude increases over days (**Figure 2A**). The VEP on day 6 is significantly larger than day 1 (**Figure 2B**, *median peak-to-peak difference: 285.05 µV, 99% CI = [283.92 290.34] µV, n = 13 mice*). In the same mice, on day 1 the normalized spectrum displayed strong high frequency power (**Figure 2C**). As the stimuli became familiar over subsequent days, high frequency power diminished and low frequency power increased. Comparing day 6 to day 1 showed day 6 had significantly more low frequency power (**Figure 2D**, *peak: 15.50 Hz, median difference: 2.88 dB, 99% CI = [1.99 3.67] dB, n = 13*) and less high frequency power (**Figure 2D**, *peak: 64.82 Hz, median difference: -1.90 dB, 99% CI = [-3.17 -0.53] dB, n = 13).* Thus, stimulus familiarity increases low frequency power and decreases high frequency power in layer 4 of V1.

The change in average spectrum power could be a result of a blanket increase in any given frequency at all times, more periods of sustained oscillatory activity, or some combination of both. We therefore measured the amount of time spent by the LFP within each frequency on each day. For our purposes, this was achieved with P-Episode (see Methods), a technique that only counts an oscillation as active if it surpasses both a power and duration threshold (Caplan et al., 2001; van Vugt et al., 2007). On day 1, the LFP spent more time exhibiting high frequency oscillations than low (**Figure 2E**). By day 6, time spent in high frequency oscillations dropped and time spent in low frequency oscillations increased. Comparing day 6 to day 1 revealed that more time was spent in low frequency oscillations on day 6 than day 1 (**Figure 2F**, *peak: 15.00 Hz, median difference: 15.24%, 99% CI = [10.81 20.39]%, n = 13*) and less time was spent in high frequency oscillations on day 6 than day 1 (**Figure 2F**, *peak: 65.00 Hz, median difference: -9.64%, 99% CI = [-16.42 -3.61]%, n = 13).* Thus, experience with a stimulus increases time spent in low frequency oscillations and decreases time spent in high frequency oscillations.

### Experience-dependent oscillations in V1 are stimulus-specific

We next sought to determine if, like SRP, the shift in frequency composition of the LFP was stimulus specific. On day 7, in addition to the now highly familiar stimulus orientation, we presented a novel stimulus that was offset by 90° from familiar. Five blocks of each stimulus were pseudo-randomly interleaved with each other. The familiar orientation induced a larger VEP than the novel orientation (**Figure 3A**), as has been observed in numerous previous studies (see, e.g., (Frenkel et al., 2006; Cooke et al., 2015; Fong et al., 2020)). Statistics confirm that the familiar VEP is significantly larger than the novel VEP (**Figure 3B**, *median peak-to-peak difference: 266.21 µV, 99% CI = [262.03 270.00] µV, n = 13*). Consistent with our observation of changes in frequency composition with growing familiarity (**Figure 2**), the familiar stimulus generated more low frequency power and less high frequency power in the layer 4 LFP than the novel stimulus (**Figure 3C**). Bootstrapping confirmed that the familiar orientation produced more low frequency power (**Figure 3D**, *peak: 14.65 Hz, median difference: 3.72 dB, 99% CI = [2.99 4.53] dB, n = 13*) and less high frequency power than the novel orientation (**Figure 3D**, *peak: 67.63 Hz, median difference: -1.92 dB, 99% CI = [-2.80 -1.14] dB, n = 13*). We observed similar results for the time spent in high and low frequency oscillations (**Figure 3E**), with more time spent in low frequency oscillations for the familiar orientation compared to the novel orientation (**Figure 3F**, *peak: 15.00 Hz, median difference: 16.25%, 99% CI = [11.76 21.12]%, n = 13*) and less time spent in high frequency oscillations (**Figure 3F**, *peak: 69.00 Hz, median difference: -11.83%, 99% CI = [-18.88 -6.32]%, n = 13*). Thus, oscillations within V1 are experience-dependent and stimulus-specific.

### Neither movement nor arousal account for the changes in layer 4 LFP frequency composition

Studies have shown that locomotion can have a substantial effect on V1 oscillations and response properties in awake mice (Niell and Stryker, 2010; Bennett et al., 2013; Fu et al., 2014; Reimer et al., 2014; Vinck et al., 2015). Given the evidence that SRP and learned suppression of behavior both occur in tandem and require the same mechanisms (Cooke et al., 2015; Kaplan et al., 2016), we were interested to understand if oscillations that emerged in the LFP with growing stimulus familiarity were simply the result of reduced movement. Our previous analyses of behavior during SRP were restricted to the first few seconds after a transition from gray screen to the stimulus, measuring an orienting or startle response that was more likely for novel than familiar stimuli (Cooke et al., 2015; Kaplan et al., 2016; Fong et al., 2020). The oscillations under investigation here extend throughout each stimulus block, over 200 seconds, so it was critical to analyze the animal’s movement over this time period. To this end, we recorded piezo-electric activity that measured ongoing forepaw movement (**Figure 4A**). The data shown in **Figure 4** exclude the first four seconds of each block to remove the contribution of an orienting or startle response but including these measurements in the average did not change the results. Analysis of the average forepaw movement, normalized to gray screen (see Methods), revealed no difference if the mice viewed familiar or novel stimuli (**Figure 4B**, *n = 11*). This was confirmed with non-parametric hierarchical bootstrapping (**Figure 4C**, *the confidence interval included 0, n = 11*). At no point within a phase reversal did novel stimuli elicit more forepaw movement than familiar stimuli or vice versa.

We simultaneously acquired the LFP and piezoelectric data in a subset of these animals over the same time interval. As expected from our previous results (**Figure 3**), the familiar stimulus generated more low frequency power (**Figure 4D**, *peak: 13.92 Hz, median difference: 4.27 dB, 99% CI = [2.83 5.52] dB, n = 5*) and less high frequency power in the layer 4 LFP compared to the novel stimulus (**Figure 4D**, *peak: 67.63 Hz, median difference: -2.74 dB, 99% CI = [-4.05 -1.56] dB, n = 5*). Thus, the changes in spectral activity driven by stimulus novelty cannot simply be accounted for by movement.

While movement itself may not account for the V1 oscillations that we have reported, changes in the LFP frequency composition could reflect global arousal shifts. Global arousal can be reliably monitored using pupillometry (Reimer et al., 2014; Reimer et al., 2016). Thus, we also tracked pupil dilation as mice underwent the SRP paradigm (**Figure 4E**). To remain consistent with the movement analysis, we excluded the first four seconds of each block of stimulation, but including them in the average did not change the results. As shown in **Figure 4F**, the average pupil diameter showed no observable or statistical difference between familiar and novel stimulus viewing conditions (**Figure 4G**, *the confidence interval included 0, n = 9*). Additionally, at no point within a phase reversal did novel stimuli elicit a larger pupil diameter than familiar stimuli or vice versa, nor was there an appreciable difference in average pupil position (∼4-6 pixels).

We simultaneously acquired the LFP and pupillometry data over the same time interval in a subset of these animals. As expected from our previous results (**Figure 3**), the familiar stimulus generated more low frequency power (**Figure 4H**, *peak: 14.04 Hz, median difference: 3.99 dB, 99% CI = [3.01 5.09] dB, n = 4*) and less high frequency power in the layer 4 LFP compared to the novel stimulus (**Figure 4H**, *peak: 67.02 Hz, median difference: -2.17 dB, 99% CI = [-3.18 -1.20] dB, n = 4*). Thus, the familiarity-dependent changes in spectral activity that we have described cannot simply be accounted for by a global arousal shift.

While there is no average pupil difference between familiar and novel stimuli (**Figure 4F-G**), we were interested in whether there might be a difference at the start of a stimulus block. Our data show that novel stimuli cause a slightly elevated pupil diameter the few seconds after block onset compared to familiar (**Figure 4I**, *n = 9*). However, this difference is not statistically significant (**Figure 4J**, *all confidence intervals after block onset included 0, n = 9*). Given that the one significant point occurs *before* the stimulus starts, it is likely a Type 1 error (false positive). Thus, the pronounced familiarity-dependent spectral activity that we observe in our paradigm is unlikely to be accounted for by a temporary or sustained global arousal shift.

### Oscillations and VEP magnitudes correlate

Given our previous measurements of increased VEP magnitude during SRP (Frenkel et al., 2006; Cooke and Bear, 2010; Cooke et al., 2015) and the concomitant changes in oscillations described here, we investigated the correlation between these two measures of experience-dependent plasticity. We analyzed the VEP magnitudes elicited by each phase reversal and the LFPs that immediately preceded them (see Methods). Changing the analysis window to the LFP after the VEP did not qualitatively change the results (*data not shown*). Cumulative distribution functions of all valid trials from all mice (see Methods) are shown for the max low frequency (10-30 Hz) power, the max high frequency (60-80 Hz) power, and the VEP magnitude (**Figure 5A-C**, *n = 13 mice*). They show that familiar stimulus presentations have more low frequency power (**Figure 5A**, *two-sample Kolmogorov-Smirnov test, p = 0.00*), less high frequency power (**Figure 5B**, *two-sample Kolmogorov-Smirnov test, p = 1.52 * 10^-68^*), and larger VEP magnitudes (**Figure 5C**, *two-sample Kolmogorov-Smirnov test, p = 2.89 * 10^-238^*) compared to novel stimulus presentations.

We then proceeded to correlate every combination of the three groups with each other. In **Figure 5 (D, F, H)** we show the correlations from one representative animal whose dataset, as a whole, is most similar to the population average (all animals are shown in **Figure 6**). In the exemplar, the max low frequency power negatively correlates with max high frequency power (**Figure 5D**). The population average for all animals shows a ∼40% negative correlation regardless of stimulus (**Figure 5E**, *median correlation for both: -0.41, 99% CI = [-0.51 -0.30]; median correlation for familiar: -0.35, 99% CI = [-0.43 -0.25]; median correlation for novel: -0.35, 99% CI = [-0.48 -0.20]; n = 13*). There is no statistical difference between the correlation for familiar and novel stimuli (*median correlation difference: 0.00, 99% CI = [-0.14 0.15], n = 13, data not shown*). In the exemplar, the max low frequency power positively correlates with VEP magnitude (**Figure 5F**). The population shows a ∼15-30% correlation depending on stimulus (**Figure 5G**, *median correlation for both: 0.33, 99% CI = [0.25 0.39]; median correlation for familiar: 0.14, 99% CI = [0.08 0.21]; median correlation for novel: 0.24, 99% CI = [0.12 0.33]; n = 13*). However, there is no statistical difference between the correlation for familiar and novel stimuli (*median correlation difference: -0.10, 99% CI = [-0.21 0.03], n = 13, data not shown*). Finally, in the exemplar, the max high frequency power negatively correlates with VEP magnitude (**Figure 5H**). The population shows a ∼20% negative correlation regardless of stimulus (**Figure 5I**, *median correlation for both: -0.25, 99% CI = [-0.33 -0.16]; median correlation for familiar: -0.17, 99% CI = [-0.25 -0.09]; median correlation for novel: -0.21, 99% CI = [-0.30 -0.11]; n = 13*). There is no statistical difference between the correlation for familiar and novel stimuli (*median correlation difference: 0.04, 99% CI = [-0.08 0.15], n = 13, data not shown*). Thus, VEPs and low-frequency oscillatory power correlate regardless of stimulus novelty, and both grow with increased familiarity. These data are compatible with the hypothesis that the same underlying biology is responsible for both manifestations of SRP.

### Experience-dependent differences in V1 emerge after the first presentation

A previous study showed that the initial VEPs and principal cell calcium transients elicited in layer 4 by the transition from gray screen to an oriented grating stimulus are the same for both familiar and novel stimuli (Kim et al., 2019). The robust familiar-novel differences observed in time-averaged VEPs and cellular responses emerge during the course of a block of stimulation. Examination of VEPs in the same animals we have used for LFP analysis confirmed this prior finding (**Figure 7A**). The first presentation of a stimulus after the gray period did not show a significant familiar-novel difference in VEP magnitude (**Figure 7B**, trial 1 *median peak-to-peak difference: 86.75 µV, 99% CI = [-60.02 226.49] µV, n = 13*). However, by the second presentation (corresponding to the first phase-reversal), a familiar-novel difference was seen (**Figure 7B**, trial 2 *median peak-to-peak difference: 190.76 µV, 99% CI = [33.33 386.49] µV, n = 13*). The emergence of SRP after the stimulus onset indicates recruitment of different circuits for familiar and novel stimuli (Kim et al., 2019).

These findings motivated us to compare the LFP oscillations proximal to the first and last stimulus presentations (both with the same 0° phase, called a flip). Consistent with other measures of SRP, the first flip in a familiar block produced little low frequency power but large high frequency power, whereas the last flip displayed the expected increase in low frequency power and decrease in high frequency power (**Figure 8A**). Compared to the first flip, the last flip in familiar blocks had more low frequency power (**Figure 8B**, *peak: 21.48 Hz, median difference: 3.16 dB, 99% CI = [1.35 4.92] dB, n = 13*) and less high frequency power (**Figure 8B**, *peak: 67.02 Hz, median difference: -2.72 dB, 99% CI = [-4.49 -0.76] dB, n = 13*). The change in high frequency power, but not low frequency power, was also seen for novel blocks (**Figure 8C**). Compared to the first flip, the last flip in novel blocks had roughly the same low frequency power (**Figure 8D**, *peak: 47.97 Hz, median difference: -1.80 dB, 99% CI = [-3.22 - 0.35] dB, n = 13*) and less high frequency power (**Figure 8D**, *peak: 68.73 Hz, median difference: -3.01 dB, 99% CI = [-4.92 -1.33] dB, n = 13*). Thus, prolonged exposure to a stimulus decreases high frequency power regardless of stimulus familiarity, while only familiar stimuli show an increase in low frequency power within a stimulus block.

We were next interested in exploring how quickly these modes of cortical activity emerge. At stimulus onset, the max normalized spectral power in the 10-30 Hz band quickly increased for familiar stimuli (**Figure 8E)**. Unexpectedly, power in this band also increased following novel stimulus onset, but the magnitude of the increase was less than for a familiar stimulus (**Figure 8F**). The max normalized spectral power in the 60-80 Hz band increased abruptly at the transition from black screen to gray screen, as expected (Saleem et al., 2017), and increased further upon exposure to a novel stimulus (**Figure 8G**). Power in this band decreased progressively over the first few phase reversals for both novel and familiar stimuli, but the familiar-novel difference was maintained (**Figure 8H**).

We next assessed how the spectral power continues to evolve as visual stimulation continues. To do this, we first created a concatenated spectrum (**Figure 9A**). Briefly, this concatenated spectrum is composed of the power spectrum for each phase-reversal, excluding the time period containing the VEP (see Methods). In agreement with previous results (**Figure 8**), the first few phase-reversals showed a different oscillatory signature than the last few phase-reversals. The concatenated spectrum appeared to be stable by the 25^th^ phase-reversal (50 sec. from stimulus onset). Thus, for each phase-reversal, we extracted the max power in the 10-30 Hz and 60-80 Hz frequency bands, and used the average of this extracted max power for presentations 26-100 as a comparator in our bootstrapping. For familiar stimuli, low frequency power started low then quickly increased to a steady value (**Figure 9B**). Bootstrapping confirmed that only the first presentation is different from the last 75 trials (**Figure 9C**, *first presentation, median difference: -2.54 dB, 99% CI = [-3.71 -1.10] dB, n = 13*). For novel stimuli, there were interesting onset dynamics in the low frequency band (**Figure 9D**). Max power in the 10-30 Hz band started at the average level, increased transiently, then decayed back to average (**Figure 9E**, *fifth presentation, median difference: 1.95 dB, 99% CI = [0.76 3.11] dB, n = 13*).

A similar analysis was conducted for high frequency power. For familiar stimuli, power in the high frequency band started high but quickly dropped to a steady level (**Figure 9F-G**, *first presentation, median difference: 2.03 dB, 99% CI = [0.95 3.03] dB, n = 13*). Similar kinetics were observed during novel stimulus viewing (**Figure 9H-I**, *first presentation, median difference: 2.16 dB, 99% CI = [0.89 3.37] dB, n = 13*), but both the transient and sustained power was shifted to greater values relative to familiar stimulus viewing.

### Layer 4 PV+ interneuron activity is suppressed as stimuli become familiar

Considerable evidence indicates that PV+ inhibitory neurons play a critical role in the generation of cortical oscillations at frequencies ≥ 40 Hz (Cardin et al., 2009; Korotkova et al., 2010; Carlen et al., 2012; Gonzalez-Burgos and Lewis, 2012; Lewis et al., 2012; Kuki et al., 2015; Jadi et al., 2016; Polepalli et al., 2017; Veit et al., 2017). Given our observation here that novel stimuli elicit high frequency oscillations, we performed experiments to measure the activity of layer 4 PV+ neurons over days during induction of SRP. We expressed GCaMP7 in genetically identified cortical neurons using a Cre-dependent conditional expression system and imaged cells with a 2-p microscope (**Figure 10A-B**). Only those cells that could be tracked across all days were included in the analysis. The average PV+ cell activity decreased over days as the animal became familiar with the stimulus (**Figure 10C**). Day 4 activity was significantly lower than day 1 activity (**Figure 10D**, *median difference: -0.13 dF/F, 99% CI = [-0.22 -0.04] dF/F, n = 9*), and less than gray screen baseline activity. On day 5, when both familiar and novel stimuli were presented, the average PV+ cell activity for each mouse showed a clear increase in activity during novel stimuli compared to familiar stimuli (**Figure 10E**). Non-parametric hierarchical bootstrapping confirmed that familiar stimuli elicited less activity than novel stimuli (**Figure 10F**, *median difference: -0.14 dF/F, 99% CI = [-0.18 -0.10] dF/F, n = 9*). Thus, layer 4 PV+ cells in V1 are activated by novel stimuli and suppressed by familiar stimuli.

### Layer 4 SOM+ interneuron activity grows as stimuli become familiar

There is considerable evidence that SOM+ inhibitory neurons contribute to low-frequency oscillations in the 15-30 Hz (“beta”) range and become more active with experience (Kato et al., 2015; Makino and Komiyama, 2015; Hamm and Yuste, 2016; Chen et al., 2017; Veit et al., 2017; Khan et al., 2018). As we observed a sharp increase in oscillations at this frequency with increasing stimulus familiarity, we also measured the activity of layer 4 SOM+ neurons over days as SRP was induced. As for the PV+ cells, we expressed GCaMP7 in SOM+ cortical neurons using a Cre-dependent conditional expression system and only analyzed cells that could be tracked across all days. The average SOM+ cell activity increased over days (**Figure 10G**). Day 4 activity was significantly higher than day 1 activity (**Figure 10H**, *median difference: 0.22 dF/F, 99% CI = [0.07 0.38] dF/F, n = 7*). On day 5, SOM+ cells were much more active during familiar stimulus viewing than during novel stimulus viewing (**Figure 10I**). Bootstrapping confirms that familiar stimuli induced more activity than novel stimuli (**Figure 10J**, *median difference: 0.41 dF/F, 99% CI = [0.28 0.56] dF/F, n = 7*).

Differences in the activity of PV+ and SOM+ cells during familiar and novel stimulus viewing were robust and surprisingly uniform. In **Figure 11** we compare activity for each neuron on days 1 and 4 as an initially novel stimulus becomes familiar, and the activity to this now-familiar stimulus to a novel orientation (**Figure 11A-B**, n = 1,251 PV+ neurons from 9 mice; **Figure 11C-D**, n = 1,021 SOM+ neurons from 7 mice). As this analysis shows, with very few exceptions, activity of PV+ cells is higher to a novel stimulus than to a familiar stimulus. Conversely, virtually the entire network of SOM+ neurons in layer 4 is more active when a familiar stimulus is viewed than when a novel stimulus is presented.

### Experience-dependent differences in PV+ and SOM+ cell activity in V1 emerge over presentations

As with our study of the oscillatory power at block onset (**Figure 8**), we sought to better understand how PV+ and SOM+ neurons in layer 4 change over the initial portion of visual stimulation. For both familiar and novel stimuli, PV+ cell activity increased rapidly upon the transition from gray to grating, and then diminished as the stimulus phase reversed (**Figure 12A**). However the PV+ cell activity significantly differed between familiar and novel stimulus conditions less than a second from block onset (**Figure 12B**, *0.70 seconds, median difference: -0.082 dF/F, 99% CI = [-0.158 -0.012] dF/F, n = 9*).

SOM+ cell activity also increased after the transition from gray screen to grating for both familiar and novel stimuli (**Figure 12C**). However, when the grating orientation was familiar, this increase occurred more rapidly than when the orientation was novel. The responses were clearly different after 8 seconds of stimulation (*8.09 seconds, median difference: 0.117 dF/F, 99% CI = [0.002 0.334] dF/F, n = 7*), and first became statistically different within 4 seconds of block onset (**Figure 12D**, *3.52 seconds, median difference: 0.055 dF/F, 99% CI = [0.001 0.128] dF/F, n = 7*).

## Discussion

A considerable body of evidence suggests that *induction* of SRP requires mechanisms that are shared with the phenomenon of LTP at excitatory synapses on principal neurons (Frenkel et al., 2006; Cooke and Bear, 2010; Aton et al., 2014; Cooke and Bear, 2014; Cooke et al., 2015). Here we examined the hypothesis that *expression* of SRP depends on the differential recruitment of inhibitory networks by familiar and novel visual stimuli. Our results show that novel stimuli activate a population of PV+ interneurons and elicit an increase in the power of high frequency oscillations in the layer 4 LFP. Across days, as a stimulus becomes familiar, PV+ cell activity and high-frequency oscillations subside, while SOM+ cell activity and low-frequency oscillations increase. Like other manifestations of SRP, these changes in LFP oscillations and interneuron activity are not subtle—they reflect dramatic shifts in the mode of visual information processing as a visual stimulus becomes familiar over days. Although the electrophysiological signature of stimulus recognition is not expressed immediately on the transition from a gray screen to a familiar stimulus (Kim et al., 2019), it does emerge quickly as evidenced by the rapid increase in low-frequency LFP power and VEP amplitude.

These observations inform and constrain the potential mechanisms that give rise to SRP. Although the current study was not designed to measure visual recognition behaviorally, extensive previous work has shown that SRP is a reliable biomarker of the changes in V1 that accompany formation, expression, and maintenance of visual recognition memory (Cooke et al., 2015; Kaplan et al., 2016; Fong et al., 2020). The changes reported by the LFP and VEPs occur over a time-course that appears to be sufficiently fast to account for recognition measured behaviorally in this assay (Cooke et al., 2015).

### Differential recruitment of mutually interacting networks of inhibitory neurons herald novelty detection and familiarity recognition

The first exposure of a mouse to an unexpected visual stimulus triggers a rapid increase in high frequency LFP and PV+ cell activity in layer 4 of V1 that continues throughout the entire block of stimulation. This is the dominant processing mode for novel visual stimuli in V1 of awake mice. One day later, when the stimulus is no longer novel, the PV+ neurons cease to respond strongly. By the 4^th^ day, presentation of the now-familiar stimulus instead causes suppression of a substantial fraction of PV+ neurons in layer 4 and, unsurprisingly, there is a clear decrease in the power and duration of high frequency oscillations of the LFP. Over the same time course, there is a substantial increase in the magnitude of the VEP elicited by the familiar stimulus. These observations are consistent with previous findings that silencing of PV+ neurons locally within V1 causes a decrease in 60-80 Hz LFP power (Chen et al., 2017; Veit et al., 2017) and an increase in VEPs that mimics and occludes SRP (Kaplan et al., 2016). Conversely, it has been shown that optogenetic stimulation of the PV+ neurons reverses SRP expression in the VEP (Kaplan et al., 2016). Together, these observational and interventional data suggest that expression of SRP in the VEP may be accounted for entirely by differential recruitment of PV+ interneurons by familiar and novel visual stimuli.

In layer 4 of sensory cortex, PV+ inhibitory neurons are known to be strongly inhibited by SOM+ neurons (Pfeffer et al., 2013; Xu et al., 2013). Inspired by a study in auditory cortex showing that passive sound exposure upregulates SOM+ neuron activity in layer 3 (Kato et al., 2015), we examined the effect of visual grating familiarity on the activity of SOM+ neurons in layer 4 of V1. The data show a robust and strikingly uniform increase in the activity of SOM+ cells as the stimulus becomes familiar. As expected from previous work (Chen et al., 2017; Veit et al., 2017), engagement of SOM+ cells by the familiar stimulus was associated with an increase in the power and duration of low-frequency LFP oscillations. This is the dominant processing mode for familiar stimuli in V1 of awake mice.

It has been shown by others that the activity of SOM+ and PV+ neurons of mouse V1 is strongly modulated by locomotion (Fu et al., 2014). Our previous studies have shown that reflexive forepaw movements (vidgets) occur for the first few seconds following the transition from a gray screen to a novel grating, and that this response diminishes over days as the grating becomes familiar (Cooke et al., 2015; Kaplan et al., 2016; Fong et al., 2020). However, using this same approach to monitor continuous forepaw movement over the entire 3.5-minute block of phase-reversing stimuli, we observed no familiar-novel differences. This finding suggests that movement is not a confounding variable for the interpretation of our LFP or imaging data collected over the same time period. Moreover, both populations of interneurons in layer 4 show comparable increases when movement occurs during visual stimulation (Pakan et al., 2016).

Thus, the differential recruitment of these inhibitory networks by familiar and novel stimuli is unlikely to be accounted for by movement.

Cortical responsiveness and oscillations are influenced by transitions in global brain states, mediated by diffusely projecting neuromodulatory systems (Hasenstaub et al., 2007; Bennett et al., 2013; Luczak et al., 2013; Kissinger et al., 2018). Our findings could be explained if novel stimuli produce more sustained arousal than familiar stimuli. However, we monitored global arousal through pupillometry (Reimer et al., 2014; Vinck et al., 2015) and found no differences in pupil size during familiar or novel stimuli. Furthermore, an expression mechanism based on slowly conducting modulatory systems seems unlikely considering (1) the speed of the transition in the LFP that heralds familiarity recognition and (2) the fact that the essential synaptic modifications underlying SRP reside within V1.

### Putting the pieces together

The current study adds important new pieces to the puzzle of SRP and, by extension, visual recognition memory in V1. The original description of SRP was the robust increase in the magnitude of the VEP elicited by phase-reversing a familiar stimulus (Frenkel et al., 2006), reflecting a net increase in positive current flowing into (most likely) radially oriented apical dendrites (Cooke et al., 2015). Combined with our previous findings (Kaplan et al., 2016), the current results indicate that the simplest explanation for this increase in net current flow is reduced PV+ mediated inhibition. Given the known connectivity of SOM+ cells and their involvement in the generation of 10-30 Hz (alpha/beta) oscillations in the LFP (Kato et al., 2015; Makino and Komiyama, 2015; Hamm and Yuste, 2016; Veit et al., 2017), an appealing hypothesis is that the experience-dependent increase in the activation of SOM+ neurons by familiar stimuli accounts for suppression of PV+ neurons and potentiation of VEPs. This simple model is challenged somewhat by our imaging experiments suggesting that the activity of the entire population of SOM+ cells in layer 4 is relatively slow to discriminate familiar and novel stimuli following block onset (**Figure 12C-D**). However, it may be that only a threshold number of SOM+ neurons need to be recruited to suppress the PV+ neurons at the earliest time points. In addition, the differential response kinetics of SOM+ cells may be underestimated as a consequence of the poor temporal resolution of calcium imaging methods. Indeed, if 10-30 Hz oscillations report recruitment of SOM+ cells in V1, then the activity of these neurons increases amply fast to account for VEP potentiation as soon as it can be detected (**Figure 8E-F**). Regardless, testing this model will require direct manipulation of SOM+ cell activity in future studies.

A full description of SRP must also account for the additional observations that, when measured with calcium imaging, layer 4 principal cell activity is reduced by familiarity (Kim et al., 2019). These calcium signals reflect changes in sustained activity, as they do not report augmented peak firing rates that occur with each familiar phase reversal (Aton et al., 2014; Cooke et al., 2015; Clawson et al., 2018), but they do mirror the habituation of behavioral responses (Cooke and Bear, 2015). It is tempting to speculate that recruitment of SOM+ neurons by familiar stimuli could be responsible for multiple facets of SRP: suppression of both principal cell and PV+ activity in layer 4, as well as the behavioral response. We are still left with the question of how SOM+ neurons become more active as a stimulus is learned. There are many possibilities that remain to be explored, but available data indicate there is an essential role for mechanisms of excitatory synaptic plasticity (Cooke and Bear, 2010, 2014).

The data suggest that the PV+ neurons, which are known to receive a more powerful thalamic input than glutamatergic principal neurons (Cruikshank et al., 2007), are highly engaged by unexpected feedforward sensory input, while SOM+ neurons are recruited by a recognition memory trace within V1 (Frenkel et al., 2006; Cooke and Bear, 2010; Cooke et al., 2015). Thus, the novelty response may reflect a default feedforward, plasticity-promoting state that persists until a stimulus is recognized as familiar. This putative organization is similar conceptually to the comparator model of habituation (Sokolov, 1963), in which sensory input is constantly compared to engrams distributed throughout the cortex and only when a match occurs is inhibition recruited to suppress reflexive behavioral output. Future studies aimed at dissecting the interplay between these two inhibitory neuronal populations within the framework of comparator/adaptive filtration systems will be of great interest.

## Acknowledgements

We acknowledge the invaluable support of Arnold Heynen, Nina Palisano, Jessica Buckey, Athene Wilson-Glover, Kiki Chu, and Erin Hickey. We particularly thank Dr. Robert W. Komorowski for initiating and encouraging this project. Support was provided by the National Institutes of Health (R01EY023037), the Picower Institute Innovation Fund, the Picower Young Faculty Support Fund (RWK), and a National Science Foundation Graduate Research Fellowship (DJH). SFC is funded by the Wellcome Trust (207727/Z/17/Z) and the Biotechnology and Biological Sciences Research Council (BB/S008276/1).

## Notes

### Competing Interest Statement

The authors have declared no competing interest.

